# Replication protein A binds RNA and promotes R-loop formation

**DOI:** 10.1101/2020.02.11.943977

**Authors:** Olga M. Mazina, Srinivas Somarowthu, Lyudmila Y. Kadyrova, Andrey G. Baranovskiy, Tahir H. Tahirov, Farid A. Kadyrov, Alexander V. Mazin

## Abstract

Replication protein A (RPA), a major eukaryotic ssDNA-binding protein, is essential for all metabolic processes that involve ssDNA including DNA replication, repair, and damage signaling. Surprisingly, we found here that RPA binds RNA *in vitro* with high affinity. Using native RIP method, we isolated RNA-RPA complexes from human cells. Furthermore, RPA promotes R-loop formation between RNA and homologous dsDNA. R-loops, the three-stranded nucleic acid structure consisting of an RNA-DNA hybrid and the displaced ssDNA strand, are common in human genome. R-loops may play an important role in transcription-coupled homologous recombination and DNA replication restart. We reconstituted the process of replication restart *in vitro* using RPA-generated R-loops and human DNA polymerases. These findings indicate that RPA may play a role in RNA metabolism and suggest a mechanism of genome maintenance that depends on RPA and RNA.

## INTRODUCTION

Replication Protein A (RPA) is a major ssDNA binding protein in eukaryotes (Haring et al., 2008). It is a highly conserved protein composed of three subunits, RPA70, RPA32, and RPA14, which all are essential for cell viability (Wold, 1997). RPA plays a critical role in most, if not all, metabolism processes that involve ssDNA including DNA replication, repair, transcription, and DNA damage signaling (Borgstahl et al., 2014; Chen and Wold, 2014; Wold, 1997; Zou et al., 2006). RPA binding protects ssDNA from degradation and unfolds DNA secondary structures. RPA interacts with various cellular proteins helping to coordinate DNA metabolic processes.

Recently, it was found that RPA is closely associated with R-loops *in vivo* (Nguyen et al., 2017; Wei et al., 2015; Yasuhara et al., 2018). Originally thought to be rare byproducts of transcription, R-loops are now known to form across the genomes of bacteria, yeast, and higher eukaryotes throughout the cell cycle (Lang et al., 2017; Stork et al., 2016; Tresini et al., 2015). In humans, R-loops occur over tens of thousands of genomic loci covering up to 5% of genome (Chedin, 2016; Sanz et al., 2016).

It was suggested that R-loops may play an important role during DNA repair by initiating transcription-coupled homologous recombination (TC-HR) in actively transcribed genome regions (Marnef et al., 2017; Wei et al., 2016; Yasuhara et al., 2018). It was also proposed that R-loops may promote restart of replication forks stalled at damaged DNA (Kogoma, 1997; Zaitsev and Kowalczykowski, 2000). The role of R-loops in priming replication was actually the first biological function proposed for this structure in bacteria (Itoh and Tomizawa, 1980). More recently, it was found that in eukaryotes persistent RNA-DNA hybrids initiate DNA synthesis in ribosomal DNA in a replication origin-independent manner (Stuckey et al., 2015). Being an important regulator of cellular processes such as transcription, gene expression, DNA replication, and DNA-repair, R-loops also represent a source of genome instability, if not timely processed or repaired (Aguilera and Garcia-Muse, 2012; Santos-Pereira and Aguilera, 2015). The mechanism of R-loop formation *in vivo* remains to be understood.

RPA has a strong binding affinity to single-stranded DNA (ssDNA) (Pokhrel et al., 2019; Wold, 1997), therefore it was thought that RPA association with R-loops is due to its binding to the displaced ssDNA strand generated during R-loop formation. Surprisingly, RPA binding to RNA has not been explored. It was presumed that RPA binds to RNA weakly, because in early studies the affinity of RPA for both RNA and dsDNA was estimated to be at least three orders of magnitude lower than for ssDNA (Kim et al., 1992).

However, our current data indicate that RPA binds to RNA much stronger than it was previously anticipated. We found that RPA binds RNA with high affinity (K_D_ ≈ 100 pM). Furthermore, we demonstrate that RPA has a unique ability to form bona fide R-loops by promoting invasion of RNA into homologous covalently closed duplex DNA. Using RPA-generated R-loops we reconstituted DNA synthesis *in vitro* using human DNA polymerases supporting the role of R-loops in the mechanism of DNA replication restart.

## RESULTS

### RPA binds to RNA with high affinity

First, using electrophoretic mobility shift assay (EMSA) we examined the RPA affinity for RNA. Previously, it was reported that the RPA binding affinity for RNA is approximately the same as for dsDNA and ~1000-fold lower than for ssDNA (Kim et al., 1992). Surprisingly, we found strong RPA binding to a 48 nt RNA (no. 501; Table S1) (K_D_ = 101.4 ± 17.0 pM), which is 300-400-fold higher than for homologous 48 bp dsDNA (nos.211/212) (K_D_ = 35.5 ± 7.0 nM) and only 30-60-fold lower than for a 48-mer ssDNA of the identical sequence (no.211) (K_D_ = 3.1 ± 0.6 pM) and (Figure 1; Figure S1). The presence of 100 mM NaCl, the condition that were used in previous studies, had no significant effect on the RPA affinity (K_D_=72.0 ± 10.2 pM) for RNA (no.501) (Figure S2). We also tested the RPA binding affinity for RNA-DNA hybrid (nos.501/212), which appeared to be twice lower (K_D_=85.9± 4.5 nM) than for dsDNA of identical sequence (nos.211/212) (Figure 1E; Figure S1C).

**Figure 1.**
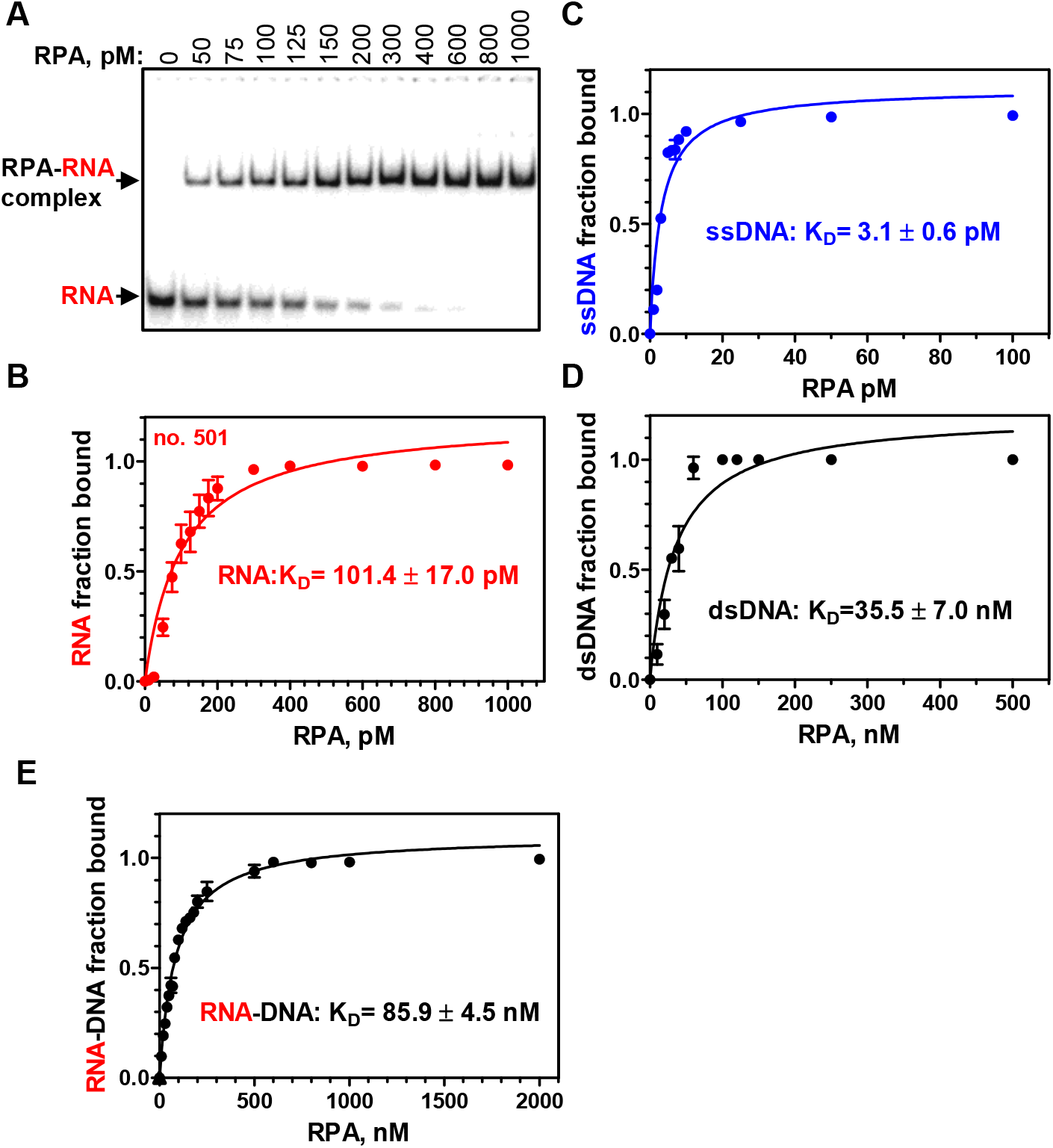
RPA binds to RNA with a high affinity. **A**, Analysis of RPA binding to a 48-mer RNA (no. 501; 5 pM) using EMSA in a 6% polyacrylamide gel. **B**, Data from (**A**) plotted as a graph. **C-E**, Graphical representation of RPA binding to 48-mer substrates: ssDNA (no.211; 0.5 pM), dsDNA (no.211/212; 3 nM), and RNA-DNA hybrid (no.501/212, 3 nM). The error bars indicate standard error of the mean (S.E.).

Then, we examined RPA binding to RNA using competitors. When RNA was used as a competitor against ssDNA, we found that the RPA affinity for RNA (no.501) is ~60-fold lower than for ssDNA of identical sequence (no.211) (Figure S3A, B). When non-homologous supercoiled pHSG299 plasmid dsDNA was used as a competitor, the affinities of RPA for RNA and ssDNA were ~500- and ~33,000-fold, respectively, higher than for plasmid dsDNA (Figure S3C, D). Thus, these results were consistent with the RPA K_D_ values for RNA and DNA indicated above.

Then, we tested the RPA binding to four other 48 nt RNAs of different sequences (Figure S4). For three of them (no.3R, no.7R, no.8R), the RPA binding affinity was high (K_D_ in the range 62.9-248.1 pM), for one of them (no.540), it was significantly lower (K_D_ > 4 nM). Inspection of the RNA structures showed that no. 540 has a much stronger propensity to form secondary structures than other tested RNAs (Table S2). Overall, these data show that RPA binds RNA with high affinity and indicate that binding is sensitive to RNA secondary structures. Several factors might contribute to underestimation of the RPA affinity for RNA in early studies including unavailability of sufficiently sensitive methods to measure RPA-RNA binding.

### RPA promotes R-loop formation *in vitro*

The finding that RPA binds RNA strongly taken together with the known association of RPA with R-loops *in vivo* prompted us to test whether RPA has the R-loop formation activity (Figure 2A). Indeed, we found that RPA can promote R-loop formation between a ^32^P-labeled 48-mer RNA (no. 501) and homologous supercoiled pUC19 plasmid DNA (Figure 2B, C). In these experiments, the plasmid DNA was prepared by a non-denaturing method to avoid formation of irreversibly denatured plasmid DNA, a source of a potential artifact due to RNA/DNA annealing. We then tested the authenticity of the RPA-promoted R-loops. In contrast to RNA-DNA hybrids produced by annealing or RNA-protein complexes that can resist deproteinization, R-loops, similar to D-loops, are sensitive to plasmid dsDNA cleavage (outside of the R-loop region) with a restriction endonuclease (Bugreev et al., 2007). The cleavage causes loss of plasmid dsDNA superhelicity and R-loop dissociation due to DNA branch migration. We found that dsDNA linearization with EcoRI indeed causes R-loops dissociation confirming their bona fide nature (Figure 2D). As expected, the R-loops were also sensitive to RNaseH, which digests RNA moiety in RNA-DNA hybrid (Figure 2E).

**Figure 2.**
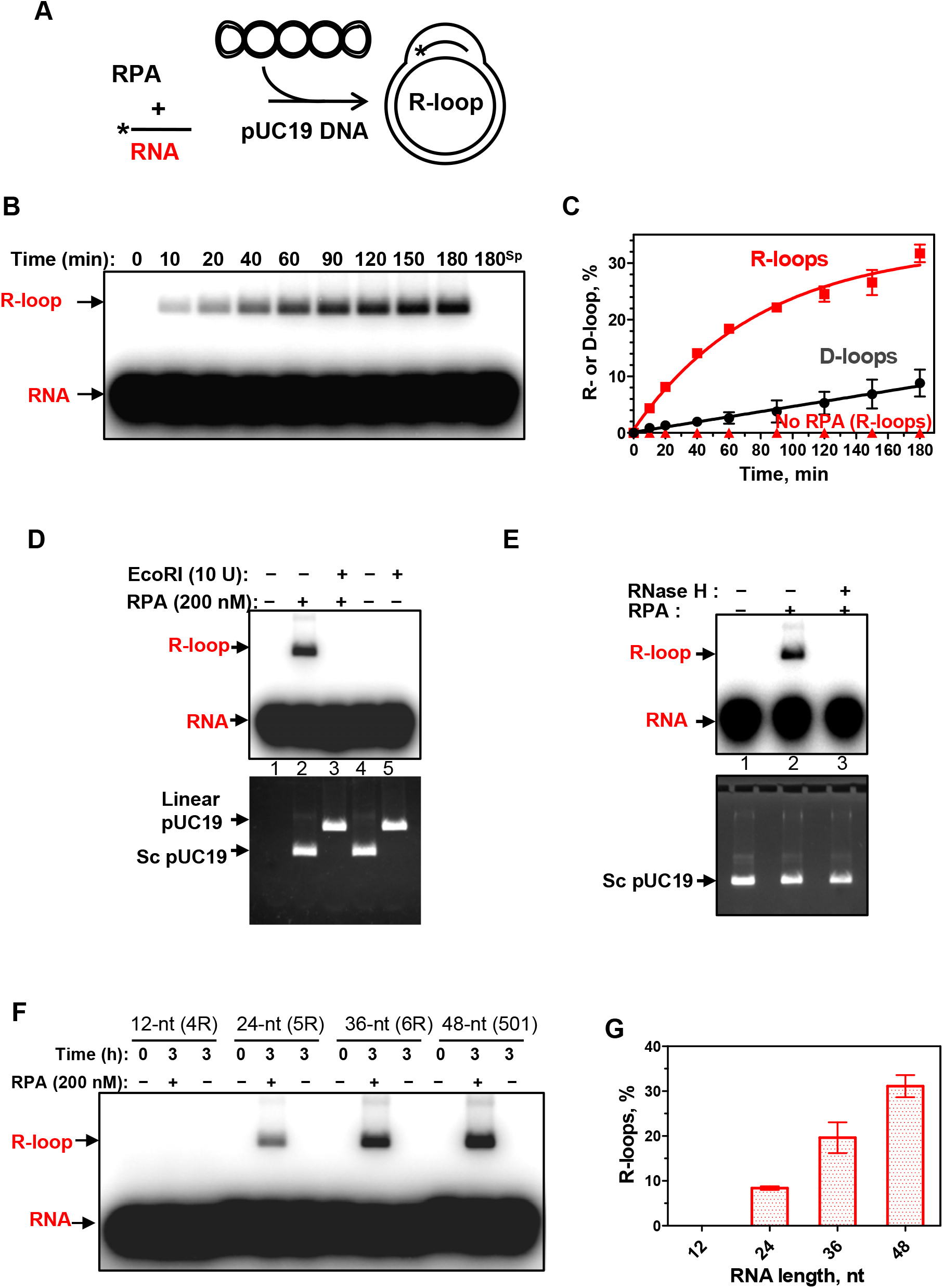
RPA promotes R-loop formation. **A**, Reaction scheme. Asterisk denotes the 32P-label. **B**, The kinetics of the R-loop formation by RPA (200 nM) analyzed by electrophoresis in a 1% agarose gel. “180^sp^” denotes the reaction (180 min) in the absence of RPA. **C**, Graphical representation of R- and D-loop formation by RPA. **D**, Sensitivity of R-loops to EcoRI. ^32^P-labeled RNA (no. 501; 3 μM, nt) was incubated with supercoiled pUC19 dsDNA (67.2 μM, nt) for 3 h in the absence (lane 1) or presence of RPA (200 nM). The R-loops were then incubated with EcoRI (lane 3). The products were analyzed by electrophoresis in 1% agarose gel. Controls include ^32^P-RNA (no. 501; 3 µM, nt) (lane 1), a mixture of ^32^P-labeled RNA (no. 501; 3 µM, nt) and pUC19 incubated with EcoRI storage buffer (lane 4) or with EcoRI (lane 5). The gel was autoradiographed (top panel) to visualize ^32^P-labeled RNA and R-loops, and then stained with ethidium bromide (bottom panel) to monitor intactness of pUC19 dsDNA. Note, R-loops co-migrate in the gel with supercoiled pUC19 DNA. **E**, Sensitivity of R-loops to RNAse H. The R-loops produced as in D (lane 2) were incubated with RNase H (5 units) (lane 3) or with the storage buffer (lane 2). The products were analyzed as in D. (**F**) RNA length-dependence of R-loop formation by RPA. R-loops were formed by RPA (200 nM) in pUC19 DNA (67.2 µM, nt) using ^32^P-labeled RNAs: 12-mer (no. 4R), 24-mer (no. 5R), 36-mer (no. 6R), and 48-mer (no. 501), (each 3 μM, nt). R-loops were analyzed by electrophoresis in a 1% agarose gel. **G**, The data from (F) presented as a graph. The error bars indicate S.E.

When RNA was replaced with a 48-mer ssDNA of the identical sequence (no.211), the efficiency of the reaction (D-loop formation) was reduced significantly indicating that the RPA activity was specific for R-loop formation (Figure 2C; Figure S5). Not all tested RNAs were equally proficient in RPA-promoted R-loop formation (Figure S6). This proficiency inversely correlates with the RNA propensity to form secondary structures (Table S2). It does not generally correlate with the RPA binding affinity for the tested RNAs, as RPA has similar K_D_ for nos. 501 and 3R that differ dramatically in their ability to support R-loop formation (Table S2). However, by titrating the RPA-RNA complexes with NaCl we found that the RPA complex with RNA no.501 is more stable than with RNA no.3R (Figure S7). Thus, the stability of RPA-RNA complexes may contribute to R-loop formation efficiency. The yield of R-loop formation rises with the increase of RNA length from 24 to 48 nt. No R-loops formed with a 12 nt RNA (Figure 2F, G) consistent with poor RPA binding to short RNAs (Figure S8). The optimal RPA concentration for R-loop formation corresponded to one RPA heterotrimer per 15 nt of RNA (Figure S9).

The R-loop forming activity shows evolutionary conservation among RPA orthologs. *S. cerevisiae* RPA *(Sc*RPA) promotes R-loop formation, albeit with an ~6-fold reduced efficiency, but the RPA functional homolog from *E. coli* (*Ec*SSB) lacks this activity under several tested conditions (Figure 3). We also found that RAD52 or RAD51 recombinase, which efficiently promoted D-loop formation, did not promote R-loop formation (Figure 4). Thus, R-loop formation appeared to be a unique activity of RPA.

**Figure 3.**
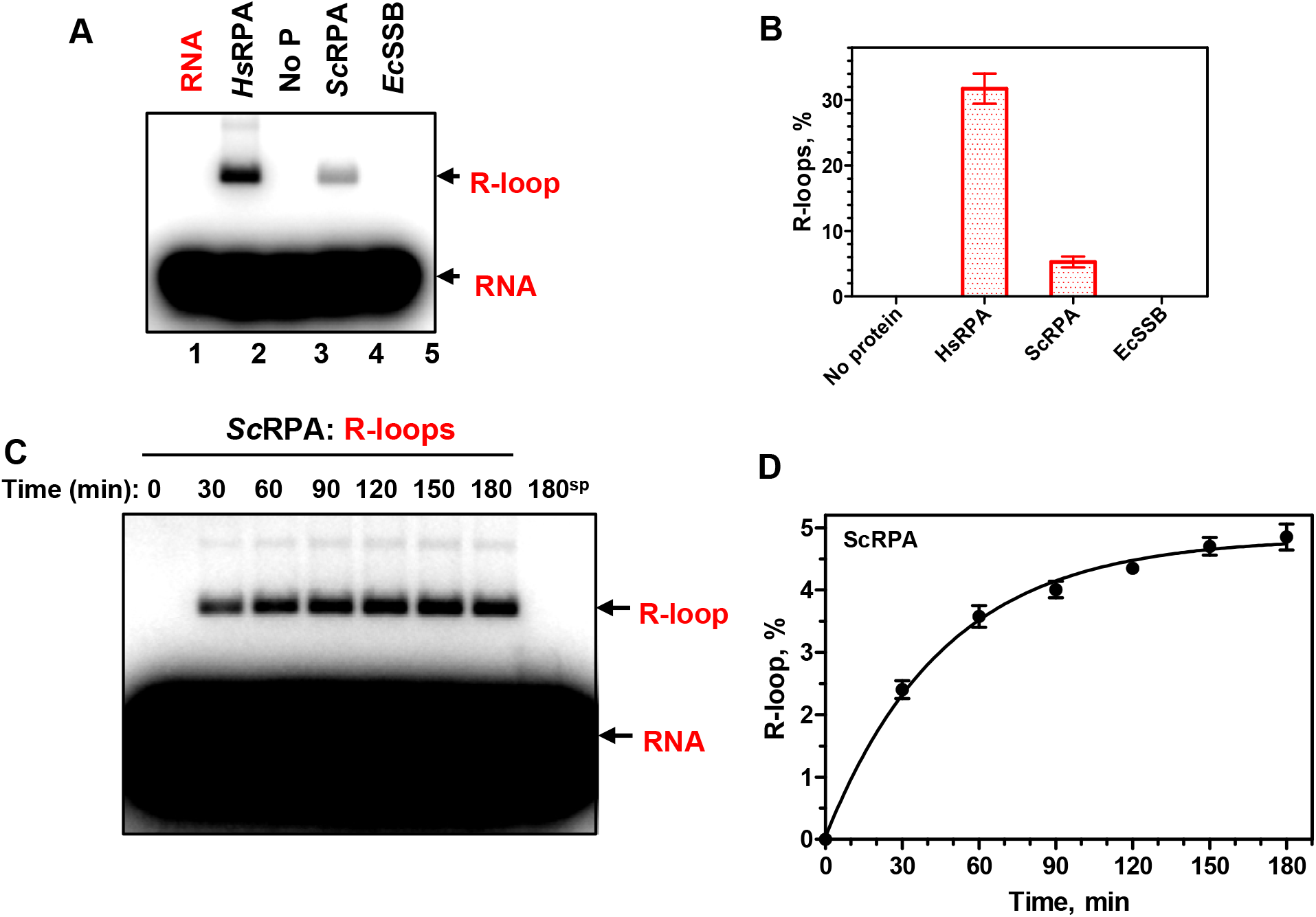
Human and yeast RPA promote R-loop formation. **A**, Human RPA (*Hs*RPA) (200 nM) and yeast RPA (*Sc*RPA) (100 nM), but not *E. coli* SSB (*Ec*SSB) (270 nM), promote R-loop formation between a 48-mer RNA (no. 501; 3 µM, nt) and pUC19 dsDNA (67.2 μM nt). The R-loops were analyzed by electrophoresis in a 1% agarose gel. **B**, The data from (A) presented as a graph. **C**, The kinetics of R-loop formation by *Sc*RPA (100 nM) between 48-mer RNA (no. 501; 3 µM, nt) and pUC19 dsDNA (67.2 μM nt) analyzed by electrophoresis in a 1% agarose gel. **D**, The data from (C) presented as a graph. The error bars indicate S.E.

**Figure 4.**
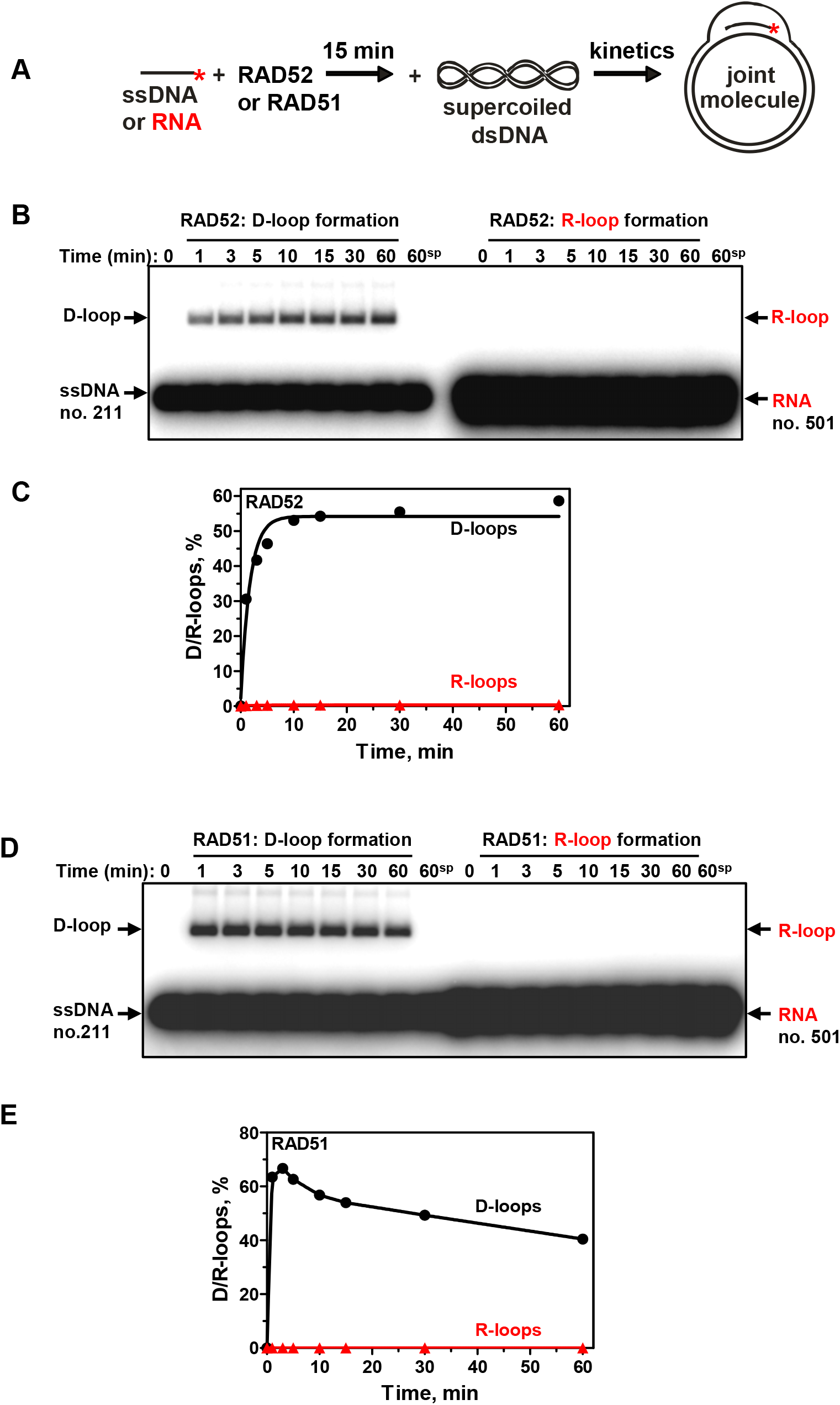
Human RAD52 and RAD51 promote formation of D-loops, but not R-loops. **A**, The scheme of D/R-loop formation. The asterisk denotes the ^32^P label. **B**, The kinetics of RAD52-promoted D- and R-loop formation. RAD52 (450 nM) was preincubated with a 48-mer ssDNA (no.211; 3 µM, nt) or RNA (no. 501; 3µM, nt) of the same sequence. The reactions were initiated by addition of supercoiled pUC19 dsDNA (50 µM, nt), the products were analyzed by electrophoresis in a 1% agarose gel. “60sp” denotes RAD52-independent (spontaneous) D/R loop formation after 60 min of reaction. **C**, The data from A shown as a graph. **D**, The kinetics of RAD51-promoted D- and R-loop formation. RAD51 (1 µM) was preincubated with a 48-mer ssDNA (no.211; 3 µM, nt) or RNA (no. 501; 3 µM, nt). The reactions were initiated by the addition of supercoiled pUC19 dsDNA (50 µM, nt). The products were analyzed by electrophoresis in a 1% agarose gel. “60sp” denotes RAD51-independent (spontaneous) D/R loop formation after 60 min of incubation. **E**, The data from D shown as a graph.

### Reconstitution of DNA replication restart using R-loops

Previously, it was suggested by Kogoma (Kogoma, 1997) that R-loops may be used to initiate the restart of DNA replication stalled at DNA damage sites. The ability of RPA to form R-loop may be especially relevant to this hypothesis because of a strong and well documented RPA association with DNA replication. Thus, RPA was initially discovered in human cell extracts as a component essential for SV40 DNA replication *in vitro* (Fairman and Stillman, 1988; Wobbe et al., 1987; Wold and Kelly, 1988). Here we wanted to test whether human DNA polymerases pol δ, pol α, pol ε and the translesion polymerase pol η can use R-loops for initiation of DNA replication. Pol η was shown to promote DNA synthesis from homologous recombination intermediates (D-loops) (McIlwraith et al., 2005). In our experiments, DNA polymerases were directly added to the R-loops generated by RPA with ^32^P-labeled RNA (no.501) and pUC19 DNA in the presence of four dNTPs (Figure 5A). RNA extension by DNA polymerases was visualized by electrophoresis in denaturing polyacrylamide gels. We found that pol η was the most efficient in utilizing the R-loop for initiation of DNA synthesis, but most of its products were short ≤ 83 nt, as could be expected for translesion DNA polymerases (Plosky and Woodgate, 2004) (Figure 5B, lane 3). Pol α and pol δ (in the presence of RFC and PCNA) were less efficient, but generated longer DNA products, ~235 nt (approximate size of the largest R-loop that can form on pUC19 supercoiled dsDNA) and even ≥1000 nt (due to the synthesis-driven strand displacement (Stith et al., 2008)). In contrast, pol ε could not efficiently use native R-loops to initiate DNA synthesis.

**Figure 5.**
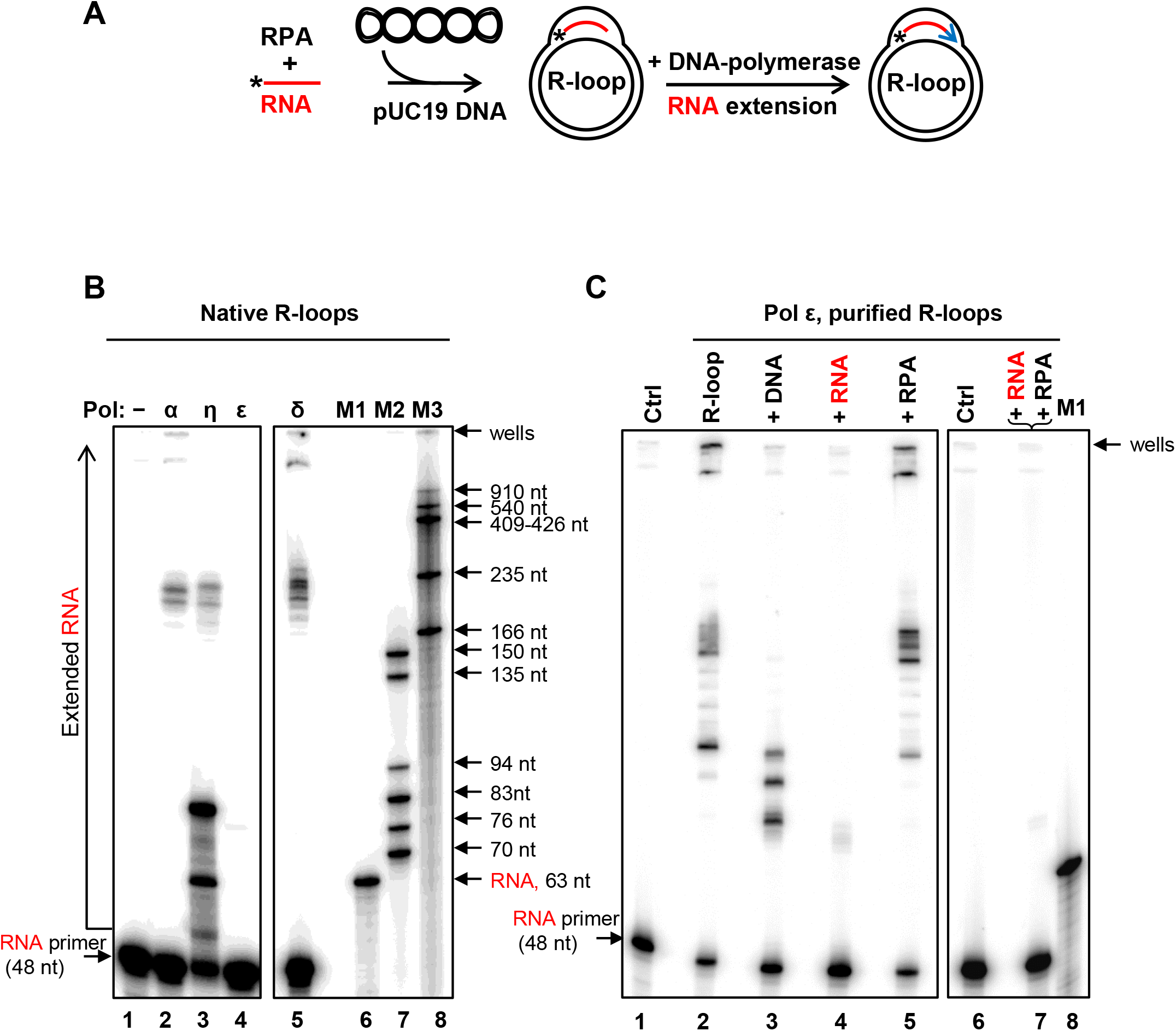
*In vitro* reconstitution of DNA synthesis restart from R-loops. **A**, Experimental scheme. Asterisk denotes the ^32^P-label at 5’-end of RNA (48 nt no.501). Blue arrow represents extension of RNA by DNA-polymerases. **B**, R-loops (3 nM) were generated in pUC19 using RPA. RNA extension in R-loops was carried out using DNA pol α (50 nM), η (38 nM), ε (50 nM), or pol δ (0.5 nM). The products of RNA extension were analyzed by electrophoresis in 8% polyacrylamide denaturing gels. In control, (lane 1) DNA polymerases were omitted. ^32^P-labeled markers shown in lanes 6-8. **C**, Effect of RNA, ssDNA, RPA and RPA-RNA on DNA synthesis by pol ε. RNA extension by pol ε (50 nM) was carried out using deproteinized and purified R-loops (1 nM) (lane 2). The R-loops were premixed with ssDNA (no.2; 3 µM, nt) (lane 3), RNA (no.517; 3 µM, nt) (lane 4), RPA (5 nM) (lane 5), or mixture of RPA (200 nM) and RNA (no.517; 3 µM, nt) (lane 7) prior to pol ε addition. In control (lane 1 and 6), pol ε was substituted with storage buffer.

Then, to evaluate the effect of RPA on the RNA extension by DNA polymerases, we deproteinized and purified R-loops. All tested DNA polymerases, including pol ε, efficiently extended RNA in the purified R-loops (Figure 5C; Figure S10). The reactions were not affected significantly when RPA was added back to purified R-loops at concentration that was sufficient to cover the displaced ssDNA strand in R-loops at stoichiometry 1 trimer per 15 nt. Surprisingly, when free RNA (no. 517; 3 µM, nt) or RPA-RNA complexes was added to R-loops we found a strong inhibition of pol ε (Figure 5C, lane 4, 7). Thus, the presence of RPA-RNA complexes inhibited activity of pol ε in the reconstitution experiments with non-deproteinized R-loops. Pol ε was also sensitive to free ssDNA (no.2; 3 µM, nt), albeit to a lesser degree (Figure 5C, lane 3). Among other tested polymerases, only pol η showed some mild sensitivity to ssDNA and RNA, and none of them showed significant sensitivity to RPA at tested concentrations (Figure S10A, B). Thus, RPA-generated R-loops can be used for initiation of DNA synthesis by human DNA polymerases: pol α, pol η, pol δ, and pol ε.

### Analysis of RPA-RNA complexes in human cells by native RIP

The observed high affinity of RPA for RNA *in vitro*, suggested that RPA may bind RNA *in vivo*. Using a native RNA immunoprecipitation (RIP) we attempted to isolate RPA-RNA complexes from the whole cell extracts (WCE) of human HEK 293T cells using mouse polyclonal anti-human RPA32 antibody (Abcam, ab88675). In preliminary experiments, we demonstrated that this antibody can recognize RPA-RNA complex, causing its supershift during electrophoresis in polyacrylamide gels (**Fig S11A**). To minimize non-specific RNA precipitation, the WCE was first pre-cleared using normal mouse IgG. We were able to isolate RPA-RNA complexes by RIP (**Fig. 6, lane 2**). The presence of RPA in these complexes was confirmed by Western blot analysis (**Fig. S11B, lane 3**). These data are in accord with the recent results of proteomics studies that identified RPA among RNA-interacting proteins in mammalian (He et al., 2016; Lee et al., 2016) and yeast cells (Mitchell et al., 2013). The RNA cross-linked peptide V_263_YYFSK_268_ was mapped in the DNA binding domain A of RPA70 (He et al., 2016). Taken together, our and others’ results show that RPA can bind RNA *in vitro* and *in vivo*.

**Figure 6.**
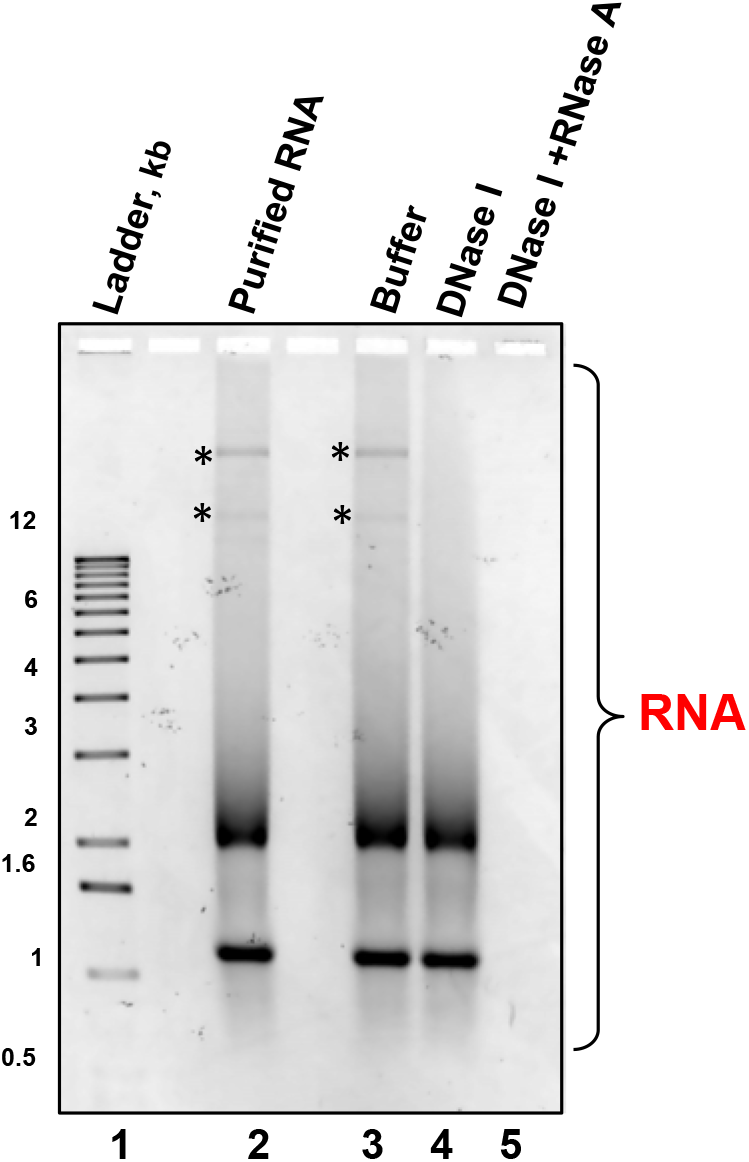
Analysis of RNA isolated from RPA-RNA complexes formed in HEK 293T cells. The RPA-RNA complexes were immunoprecipitated using mouse polyclonal anti-human RPA32 antibody from HEK 293T WCE. Equal aliquots of purified RNA (124 ng) were either left intact (lane 2), treated with DNase I (lane 4) or DNase I+ RNase A (lane 5). In control, DNase I was replaced by its storage buffer (lane 3). 1 kb DNA ladder is shown in lane 1. The samples were analyzed by electrophoresis in a 1% agarose gel. The asterisks denote traces of DNA.

Then, we purified RNA from the RIP-complexes and subjected it to new generation sequencing (Applied Biological Materials Inc). Prior to sequencing, the final preparation was confirmed to contain almost exclusively RNA (**Fig. 6, lanes 3-5**). A broad range of RNAs were uncovered by sequencing including mRNA, microRNA, small nuclear RNAs and long non-coding RNA (lncRNA) (data not shown); among lncRNAs was NORAD identified by Mendell’s group as an RPA-interacting RNA in WCE of human HCT116 cells (Lee et al., 2016).

## DISCUSSION

In this study we identified novel unanticipated activities of RPA. RPA binds to RNA with sub-nanomolar affinity, about 500-fold higher than to dsDNA and only 60-fold lower than to ssDNA. Furthermore, RPA promotes R-loop formation between RNA and homologous supercoiled dsDNA. We show that human DNA polymerases, α, δ, ε, and η, can utilize RPA-generated R-loops for initiation of DNA synthesis *in vitro* supporting a previously proposed role of R-loops in DNA replication restart (Kogoma, 1997; Zaitsev and Kowalczykowski, 2000).

The high affinity of RPA to RNA *in vitro* may suggest that RPA binds RNA also *in vivo*. Recent proteomics studies support this proposal. Thus, RPA has been identified among RNA-interacting proteins in mammalian (He et al., 2016; Lee et al., 2016) and yeast cells (Mitchell et al., 2013). Bonasio’s group by protein-RNA photo-crosslinking and quantitative mass spectrometry identified RPA among the proteins that interact with RNA regardless of its polyadenylation status in the nuclei of embryonic stem cells (He et al., 2016). The RNA cross-linked peptide V_263_YYFSK_268_ was mapped in the DNA binding domain A of RPA70. Mendel’s group identified RPA among the proteins that interact with long non-coding RNA NORAD (Lee et al., 2016). In that study, biotinylated RNA fragments of NORAD were incubated with HTT-116 whole cell lysates and the proteins that bind to these fragments were eluted and identified by mass spectrometry. RPA70, RPA 32, and RPA 14 subunits were among the proteins that specifically bind NORAD RNA. Parker’s group by UV cross-linking proteins to mRNAs identified RPA among the proteins that directly interact with mRNA *in vivo*. mRNA-protein complexes were then purified under denaturing conditions using oligo(dT) columns, and the RNA–bound proteins were analyzed by LC-MS/MS (Mitchell et al., 2013). *Sc*RFA1 subunit (ortholog of *Hs*RPA70) was identified among the mRNA-bound proteins. The biological role of RPA-RNA interactions remains to be understood. RPA may protect RNA from RNases or recruit proteins that are involved in RNA metabolism. A putative role of RPA in mRNA nuclear export was reported (Chen et al., 2012). Our native RIP data (Figure 6) are consistent with these publications. Additional studies are needed to further characterize RPA-RNA interactions *in vivo.*

Even though RNA is abundant in the cell, RPA binding to RNA may not necessarily interfere with its well-established functions in DNA metabolism that require RPA binding to ssDNA. It is likely that RPA will transfer from RNA to ssDNA generated during DNA replication stress or damage due to its 60-fold higher affinity for ssDNA. Recent studies showed a dynamic nature of RPA binding even to ssDNA, to which it has the highest affinity (Chen et al., 2016; Gibb et al., 2014; Nguyen et al., 2014; Pokhrel et al., 2019). Thus, RPA can translocate along the ssDNA axis and transfer from one polynucleotide to another.

RPA appeared to be the first know protein that promotes formation of bona fide R-loops by invading RNA into covalently closed duplex DNA. R-loops are structurally distinct from RNA-DNA hybrids that are formed either through RNA-DNA annealing or inverse RNA strand exchange occurring in the proximity of linear dsDNA ends (Keskin et al., 2014; Mazina et al., 2017; Zaitsev and Kowalczykowski, 2000). While the mechanism of R-loop formation by RPA remains to be investigated, several assumptions can be made. It is likely that during the initial step of R-loop formation, RPA acts in a complex with RNA due its ~500-fold higher affinity for RNA comparing with the plasmid dsDNA. Moreover, the optimal amount of RPA required for R-loop formation corresponds to its stoichiometric coverage of RNA, but not dsDNA (Figure S9). Next, RPA-RNA complex needs to engage dsDNA in the homology search process. The RPA trimer has six DNA binding domains (Chen and Wold, 2014; Pokhrel et al., 2019) which could potentially provide binding space to both RNA and dsDNA juxtaposing them for RNA:DNA pairing. Binding of dsDNA by the RPA-RNA complex should be by necessity week to allow multiple association-dissociation steps during the homology probing. After homology is found and initial R-loops are formed, RPA may not remain bound to the newly formed RNA-DNA heteroduplex but be transferred to the displaced ssDNA strand to which it has much higher affinity (Figure 1). This RPA binding to the displaced ssDNA strand may help to stabilize and further expand the R-loop. A relatively weaker RPA binding to RNA comparing with ssDNA may favor its R-loop formation activity as opposed to D-loop formation. Because of a strong binding to RPA ssDNA may occupy all available binding space preventing dsDNA binding that is needed for formation of D-loops.

Recent data indicate that R-loops are a common structure in genomes of humans and other species (Chedin, 2016; Sanz et al., 2016). The biological role of R-loops is currently under intense investigation. It was found that R-loops are essential for repair of DNA double-strand breaks in actively transcribed genome regions through TC-HR (Marnef et al., 2017; Wei et al., 2016; Yasuhara et al., 2018) or Non-Homologous End-Joining (Chakraborty et al., 2016). It was proposed by Kogoma that R-loops may serve as a primer to restart DNA replication stalled at DNA lesions (Kogoma, 1997) (Figure 7). However, the mechanistic support of his hypothesis was lacking as none of the known recombinases promotes R-loops formation. The R-loop formation activity of RPA may be especially relevant to replication restart because of a strong RPA association with DNA replication (Fairman and Stillman, 1988; Wobbe et al., 1987; Wold and Kelly, 1988). RPA32 subunit was directly UV crosslinked with the RNA strand of the nascent RNA-DNA primer during SV40 replication in nuclei of monkey CV-1 cells (Mass et al., 1998). It was demonstrated that RPA interacts with pol α, RFC, and pol δ (Mo et al., 2000; Yuzhakov et al., 1999). RPA stabilizes a complex between short RNA primer and pol α and then coordinates loading of RFC, PCNA, and pol δ to initiate DNA synthesis. We found that all tested human DNA polymerases, pol α, pol δ, pol ε, and pol η, can utilize RPA-generated R-loops for initiation of DNA synthesis *in vitro*. These *in vitro* reconstitution experiments further support Kogoma’s hypothesis and suggest the mechanisms of genome maintenance that depend on RPA and RNA.

**Figure 7.**
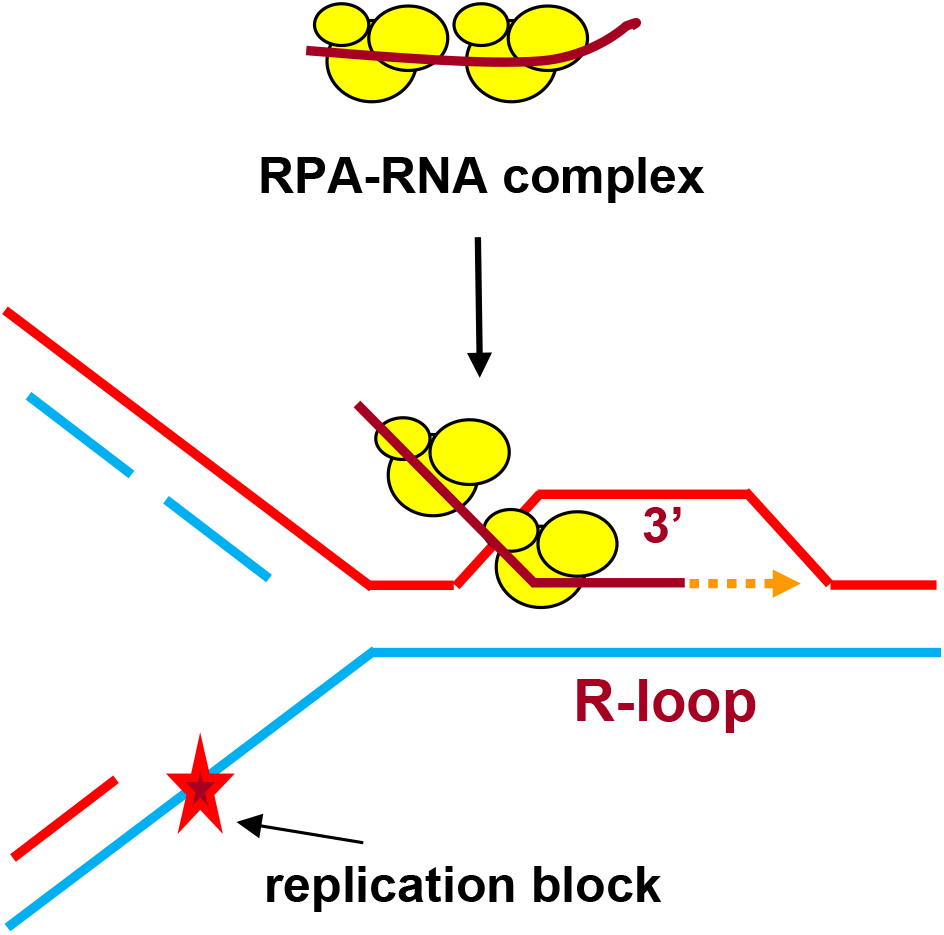
Proposed RPA-dependent DNA replication restart initiated at R-loop. RPA promotes formation of the R-loop that serves as a primer for a DNA polymerase during replication restart.

## MATERIALS AND METHODS

### Proteins, DNA and RNA

Human RPA, RAD51, RAD52, and yeast Rad52 were purified as described (Henricksen et al., 1994; Sigurdsson et al., 2001; Singleton et al., 2002; Song and Sung, 2000). Human DNA-polymerases: pol η, the catalytic core of pol α p180(335-1257), the Flag-tagged four subunit pol δ, RFC, and PCNA were purified as described (Baranovskiy et al., 2014; Kadyrov et al., 2009; Masutani et al., 2000; Rodriges Blanko et al., 2016). The catalytic core of human pol ε p261(1-1172)exo^−^, was purified according to (Zahurancik et al., 2013) with the following modifications: His-tag was placed on the N-terminus before a Sumo-tag and removed by Sumo protease after the first purification step, which included the nickel ion affinity chromatography. The oligodeoxyribonucleotides (Table S1) were purchased from IDT Inc. and further purified by electrophoresis in polyacrylamide gels containing 50% urea (Rossi et al., 2010). HPLC-purified oligoribonucleotides were purchased from IDT Inc. All experiments with RNA were carried out in the presence of 1 × Ambion RNA*secure* RNase Inactivation reagent. Double-stranded oligonucleotides were prepared by annealing of equimolar (molecules) amounts of complementary oligonucleotides (Rossi et al., 2010). When indicated, oligonucleotides were 5-end labeled with ^32^P-γATP using T4 polynucleotide kinase (New England Biolabs). Supercoiled pUC19 plasmid dsDNA was prepared by a method that did not involve DNA denaturation (Clewell and Helinski, 1969) with modifications. Briefly, *E. coli* host cells were treated with lysozyme and lysed with Triton X-100. The lysate was cleared by centrifugated at 4 °C for 30 min at 40,000 × g. The cleared lysate was mixed with ethidium bromide to 700 µg/ml and CsCl (0.59 g per 1 ml of cleared lysate) and loaded at the top of CsCl (1.58 g/ml in water) solution that filled the bottom half of the centrifuge tube. The samples were centrifuged in an angle rotor for 24 h at 200,000 × g at the ambient temperature. Isolated supercoiled plasmid DNA was further purified by 3 × butanol extractions followed by gel-filtration on a Sephacryl S-500 column. Supercoiled pHSG299 plasmid dsDNA purified by CsCl-ethidium bromide gradient centrifugation was purchased from Takara Bio Inc. pHSG299 is a derivative of pUC19 plasmid in which ampicillin resistant gene was replaced with kanamycin resistant gene. DNA and RNA concentrations are expressed in moles of molecules or, when indicated, in moles of nucleotides.

### RPA binding to RNA, ssDNA, dsDNA and RNA-DNA hybrid using EMSA

20 µl-mixtures contained human RPA at indicated concentrations, 25 mM Tris·acetate (pH 7.5), 10 mM KCl (added with the protein stock), 2 mM DTT, 1 mM magnesium acetate, 100 μg/ml BSA, and ^32^P-labeled 48-mer nucleic acid substrates: RNA (no.501; 5 pM, molecules), or ssDNA (no.211; 0.5 pM, molecules) or dsDNA (nos.211/212; 3 nM, molecules), or RNA-DNA hybrid (nos.501/212; 3 nM, molecules). The mixtures were incubated for 15 min at 37 °C, and then placed on ice. Each sample was supplemented with 3 µl of 50% glycerol and loaded onto a 6% polyacrylamide (29:1) gel in 0.25 × TBE buffer (22.5 mM Tris, 22.5 mM borate, and 0.25 mM EDTA, pH 8.3). Bromophenol blue was added only in the sample containing ^32^P-labeled probe without RPA. Electrophoresis was carried out at 13 V/cm for 1 h at room temperature. The gels were dried on Amersham Hybond-N+ membrane (GE Healthcare) and analyzed using a Typhoon FLA 7000 biomolecular imager. The K_D_ and B_max_ values were obtained by fitting the data to one site binding hyperbola in GraphPad Prism 5.0. B_max_ values were 1.20 ± 0.07, 1.11 ± 0.08, and 1.20 ± 0.07, and 1.10 ± 0.02 for RNA (no.501), ssDNA (no. 211), dsDNA (nos.211/212), and RNA-DNA hybrid (nos.501/212), respectively.

### RPA binding to RNA or ssDNA in the presence of competitors

RPA (20 pM) was incubated with a ^32^P-labeled 48-mer ssDNA (no. 211; 5 pM, molecules or 240 pM, nt) that was pre-mixed with various indicated amounts of non-labeled RNA (no. 501) or ssDNA (no.211) for 15 min at 37 °C. In other experiments, RPA at indicated concentrations was incubated with ^32^P-labeled RNA (no. 501; 5 pM, molecules or 240 pM, nt) or ^32^P-labeled ssDNA (no.211; 5 pM, molecules or 240 pM, nt) that was pre-mixed with various indicated amounts of pHSG299 supercoiled plasmid dsDNA for 15 min at 37 °C. The non-homologous pHSG299 dsDNA competitor was used to avoid possible DNA/RNA pairing that might interfere with the RPA binding measurements. RPA-^32^P-RNA and RPA-^32^P-DNA complexes were analyzed by EMSA as described above.

### D-loop and R-loop formation

Human RPA (200 nM) was incubated with ^32^P-labeled RNA (3 μM, nt) or ssDNA (3 μM, nt) in buffer A containing 25 mM Tris·acetate (pH 7.5), 10 mM KCl (added with the protein stock), 2 mM DTT, 1 mM magnesium acetate, and 100 μg/ml BSA for 15 min at 37 °C. The reactions were initiated by addition of supercoiled pUC19 dsDNA (67.2 μM, nt). Aliquots (10 µl) were withdrawn at indicated time points and deproteinized by incubation with 1% SDS, 1.6 mg/ml proteinase K, 6% glycerol and 0.01% bromophenol blue for 15 min at 37 °C. Samples were analyzed by electrophoresis in 1% agarose gels in TAE buffer (40 mM Tris·acetate, pH 8.0 and 1 mM EDTA). Electrophoresis was carried out at 5 V/cm for 1.5 h at room temperature. The gels were dried and analyzed as described above for EMSA. The D-loop/R-loop yield was expressed as a percentage of the input plasmid DNA.

For *S.* c*erevisiae (Sc)* RPA and *E. coli (Ec)* SSB, the R-loop formation was carried out as described above, except that magnesium acetate concentration was 2 mM and the protein concentrations were 100 nM and 270 nM, respectively.

For human RAD52, the R/D-loop formation reactions were carried out as described for RPA, except that RAD52 (450 nM) was used instead of RPA, magnesium acetate was 0.3 mM and supercoiled pUC19 dsDNA was 50 μM (nt). For human RAD51, the R/D-loop formation reactions were carried out as described for RPA, except that 1 mM ATP was included in the reaction mixture, 1 mM CaCl_2_ was used instead of magnesium acetate and the concentrations of RAD51 and supercoiled pUC19 dsDNA were 1 μM, and 50 μM (nt), respectively.

### Cleavage of the R-loops with EcoRI restriction endonuclease

RPA-promoted R-loop formation was carried out for 3 h at 37 °C, as described above. Then, 1.5 µl of 50 mM magnesium acetate and 0.5 µl (10 units) of EcoRI restriction endonuclease were added to 10 µl of the reaction mixture and incubation was continued for another 15 min. The samples were deproteinized and analyzed by electrophoresis in 1% agarose gels. The gels were dried and analyzed as described above for the R-loop formation. The dried gels were then rehydrated, stained with ethidium bromide, and analyzed as described above for RNase H treated R-loops.

### Treatment of the R-loops with RNase H

RPA-promoted R-loop formation was carried out for 3 h at 37 °C, as described above. Then, 1 µl of 10x RNase H reaction buffer and 1 µl (5 units) of RNase H (New England Biolabs) were added to 8 µl of the reaction mixture and incubation was continued for another 30 min. The samples were deproteinized and analyzed by electrophoresis in 1% agarose gels. The gels were dried and autoradiographed and analyzed using a Typhoon FLA 7000 biomolecular imager as described above for EMSA. The dried gels were then rehydrated by soaking in water, detached from the Amersham Hybond-N+ membrane, stained with ethidium bromide (2 µg/ml in water) for 30 min at room temperature, destained for 30 min in a large volume of water, and subjected to image analysis using an AlphaImager 3400 gel documentation station.

### Reconstitution of DNA synthesis restart using R-loops

RPA-promoted R-loop formation between pUC19 and ^32^P-labeled 48-mer RNA (no.501) was carried out in buffer containing 25 mM Tris acetate pH 7.5, 1 mM magnesium acetate, 100 µM each of four dNTPs, 250 µg/ml BSA, and 10 mM DTT for 3 h at 37 °C. Then, KCl was added to final concentration 40 mM. To initiate DNA synthesis from R-loops (3 nM, molecules) 9-µl aliquots were mixed with DNA pol α (50 nM), or pol η (38 nM), or pol ε (50 nM) and incubated for 30 min at 37 °C. Addition of the DNA– polymerases increased final KCl concentration to 60 mM.

For RNA extension by pol δ, RPA-promoted R-loop formation was performed in standard buffer A for 3 h at 37 °C. Reconstitution reactions (10 µl) contained R-loops (3nM, molecules), RFC (8 nM), PCNA (48 nM), and pol δ (0.5 nM), 30 mM Tris acetate pH 7.5, 5 mM magnesium acetate, 100 µM each of four dNTPs, 1 mM ATP, 250 µg/ml BSA, 10 mM DTT, 60 mM KCl. R-loops were pre-incubated with RCF and PCNA for 5 min at 37 °C, and then pol δ was added and incubation was continued for another 30 min.

All DNA polymerization reactions were terminated by adding 15 µl of 99.9% formamide, containing 0.1% of bromophenol blue. The samples were heated for 4 min at 80 °C, and the products of RNA extension were analyzed by electrophoresis in an 8% denaturing PAGE (19:1), containing 50% urea. The migration markers M1-M3 were 63-nt RNA, 70-150 nt ssDNA oligonucleotides and 166-910 nt denatured DdeI restriction fragments of pUC19, respectively. After electrophoresis, the gels were fixed in 10% glacial acetic acid and 10% ethanol for 20 min at room temperature, dried and analyzed using a Typhoon FLA 7000 biomolecular imager.

### RNA extension by DNA polymerases using deproteinized R-loops

RPA-promoted R-loop formation was performed in buffer A for 3 h at 37 °C. The R-loops were deproteinized by treatment with proteinase K (1 mg/ml) and 0.8% SDS for 30 min at 37 °C, and then 1 mM EDTA, pH 8.0 was added to chelate magnesium ions. The deproteinized R-loops were purified by passing twice through S-400 spin columns (GE Healthcare) equilibrated with 25 mM Tris-HCl (pH 7.5) and 25 mM KCl. The purified R-loops were supplemented with 1 mM of magnesium acetate and kept at −20 °C.

Reactions (10 µl) were initiated by adding DNA pol α (50 nM), or pol η (4 nM), or pol ε (50 nM) to deproteinized R-loops (1 nM) in 30 mM Tris-HCl (pH 7.5), 1 mM magnesium acetate, 100 µM each of four dNTPs, 250 µg/ml BSA, 10 mM DTT, 60 mM KCl and carried out for 30 min at 37 °C.

For DNA pol δ, the reactions (10 µl) were carried out in 30 mM Tris acetate pH 7.5, 5 mM magnesium acetate, 100 µM each of four dNTPs, 1 mM ATP, 250 µg/ml BSA, 10 mM DTT, and 60 mM KCl. Deproteinized R-loops (1nM, molecules) were pre-incubated with RCF (2 nM) and PCNA (10 nM) for 5 min at 37 °C, and then pol δ (0.5 nM) was added followed by incubation for another 30 min.

### Cell culture

HEK293T cells (human embryonic kidney cells, ATCC [Cat. No. CRL-3216]) were cultured in Dulbecco’s modified Eagle’s high glucose medium (Sigma, D6429), supplemented with 10% fetal bovine serum, penicillin (100 units/ml) and streptomycin (100 µg/ml). Cells were maintained at 37 °C in humidified atmosphere containing 5% CO_2_.

### Super-shift of RPA-RNA complexes using anti-RPA32 antibody

Human RPA (100 nM) was incubated with ^32^P-labeled RNA (no. 3R; 3 µM, nt) at 37 °C for 15 min in binding buffer (25 mM Tris-acetate (pH 7.5), 10 mM KCl (added with the protein stock), 2 mM DTT, 1 mM magnesium acetate, and 100 µg/ml BSA). The reaction mixture (10 µl) was then cooled on ice, mixed with 0.2 µg (in 1 µl) of primary antibody (mouse polyclonal anti-RPA32, ab88675) and incubated on ice for 1 h. Each sample was supplemented with 1.5 µl of 50% glycerol and loaded onto a 6% native polyacrylamide (29:1) gel in 0.25 × TBE buffer. Electrophoresis and image visualization were carried out as described above for EMSA.

#### Native RNA immunoprecipitation (RIP)

HEK293T cells (~ 80% confluent) were collected using trypsin/EDTA (Sigma), washed at least three times with 1 × PBS, and then lysed for 30 min on ice with buffer (25 mM Tris-HCl (pH 8.0), 120 mM NaCl, 1 mM EDTA, and 0.5% NP-40) containing 1x complete EDTA-free protease inhibitor cocktail (Roche). The cell lysate was centrifuged for 20 min at 16,000 × g at 4 °C. The supernatant (whole-cell extract or WCE) was transferred to a fresh Eppendorf test-tube on ice, and then stored at −80 °C. Protein concentration in the WCE was estimated using Bradford assay (Bio-Rad). The WCE was diluted in buffer containing 25 mM Tris-HCl (pH 8.0), 1 mM EDTA, and 1 × complete EDTA-free protease inhibitor cocktail to final protein concentration 1.5 mg/ml. To remove protein-RNA complexes, which could immunoprecipitate non-specifically, 3 µg of normal mouse IgG (protein A purified, Innovative Research) was incubated with 500 µl of diluted WCE (750 µg) for 2 h at 4 °C with gentle rotation. Then 40 µl (Santa Cruz Biotech, 25% slurry in 1 × PBS) of protein A/G plus agarose beads were added and incubation continued for another 2 h. The beads were precipitated by centrifugation at 2,500 × g for 5 min at 4 °C. The supernatant was incubated with 3 µg of mouse polyclonal primary antibody (Abcam, ab88675) raised against human RPA32 at 4 °C overnight with gentle rotation. Then, 40 µl of protein A/G plus agarose beads (25% slurry) were added to the antibody-RPA-RNA complexes, and the incubation continued for another 2 h. The beads were precipitated by centrifugation at 2,500 × g for 5 min, washed 5 times with 0.2 ml of ice-cold PBS, and then once with 25 mM Tris-HCl (pH 7.5), 150 mM NaCl, and 1 × RNA*secure* reagent (Ambion). The beads were re-suspended in the last washing buffer in a total volume of 80 µl. The agarose bead-antibody-RPA-RNA mixture (80 µl) was warmed up to 37 °C, and then deproteinized by adding of 8 µl of 10% SDS and 8 µl of proteinase K (19 mg/ml stock, Roche). Deproteinization was carried out at 37 °C for 30 min with gentle flicking. To chelate any divalent ions 2 µl of 98 mM EDTA (pH 8.0) were then added. The RNA purification was performed using RNA Clean & Concentrator-5 kit (Zymo Research) as described by manufacturer. Small amounts of contaminating DNA were removed from the RNA preparation using DNA-free kit (Ambion) as described by manufacturer.

To verify that the final preparation contains only RNA its aliquot (124 ng) was treated with RNase A (Qiagen) at concentration 25 µg/ml in buffer containing 5 mM Tris-HCl and 0.5 mM EDTA (pH 8.0) for 15 min at 37 °C in total volume of 10 µl. The RNA sample was then deproteinized by incubation in 1% SDS, 1.6 mg/ml proteinase K, 6% glycerol and 0.01% bromophenol blue for 15 min at 37 °C. The RNA samples after each purification step were analyzed in a 1% agarose gel in TAE buffer (40 mM Tris·acetate (pH 8.0) and 1 mM EDTA). The gel was stained with SYBR gold dye (Thermo Fisher Scientific) as described by manufacturer and then visualized using a Typhoon FLA 7000 biomolecular imager (GE Healthcare).

In control, the agarose bead-antibody-protein-RNA complexes were washed with PBS as described above. The beads were then re-suspended in 20 µl of PBS, mixed with 20 µl of 2 × electrophoresis sample buffer (Laemmli, 1970), and incubated for 10 min at 95°C with gentle flicking. The beads were precipitated by centrifugation at 2,500 × g for 5 min. The supernatant was then used for Western blot analysis in order to verify that the RPA protein was indeed precipitated.

### Western blot analysis

The sample of the WCE immunoprecipitated with specific anti-hRPA32 antibody was separated by 10% SDS-PAGE at 100 V (constant) and transferred to a PDVF membrane (Amersham Hybond) at 70 V (constant) for 3 h at 4 °C using the Bio-Rad mini protean apparatus. The transfer was carried out in a transferring buffer (25 mM Tris, 192 mM glycine (pH 8.3), and 20% methanol). Membranes were blocked with 5% nonfat milk in TBST buffer (10 mM Tris-HCl (pH 8.0), 150 mM NaCl, and 0.1% Tween-20) for 1 h at room temperature. Then the membranes were incubated with primary mouse polyclonal antibody against human RPA32 (Abcam, ab88675) at concentration of 375 ng/ml in TBST buffer with 5% nonfat milk at 4 °C overnight. The WCE immunoprecipitated with normal mouse IgG (protein A purified, Innovative Research) was used as a negative control. After extensive washing with TBST buffer the membranes were incubated with horseradish peroxidase-conjugated goat anti-mouse secondary antibody (Jackson Immunoresearch Laboratories Inc.) at 1:10,000 dilution in TBST buffer with 5% nonfat milk for 2 h at room temperature. Blots were developed using SuperSignal West Pico detection kit (Pierce) as described by manufacturer.

### RNA sequencing and data analysis

RNA sequencing was performed by Applied Biological Materials Inc (Canada). The quality of the RNA was assessed by Qubit RNA assay and Agilent Bioanalyzer. Sequencing library was prepared using Illumna TruSeq stranded mRNA library preparation kit. Such a library preferentially captures polyadenylated forms of both coding and non-coding RNAs. Sequencing was performed on the Illumna NextSeq platform. At least 20 million 75-nt single- or paired-end reads were obtained. We removed the sequencing adapters and low-quality reads using Trimmomatic software (Bolger et al., 2014). Good quality reads were aligned to the human genome using the software package STAR (v2.5) (Dobin et al., 2013). The reference human genome (GRCh38) and the reference General Transfer Format (GTF) file for gene annotation were downloaded from the Ensembl database (Kersey et al., 2016). The aligned output file is sorted using SAMtools (v1.3.1) (Li et al., 2009). Transcript assembly and abundance estimation was performed using cufflinks (v2.2.1) with default parameters (Trapnell et al., 2012).

## QUANTIFICATION AND STATISTISTICAL ANALYSIS

For statistical analysis, GraphPad Prism 5 software was used. *In vitro* experiments were repeated at least three times; standard errors (SE) are presented on the graphs.

## AUTHOR CONTRIBUTIONS

O.M. conducted the experiments. S.S. performed computational analysis of the RNA sequences and free energies. L.K., A.B., T.T. and F.K. purified human DNA polymerases, PCNA and RFC and help to design the DNA replication experiments. A.M. and O.M. designed and analyzed experiments, conducted all statistical analysis wrote the manuscript with input and suggestions from all authors.

## ACKNOWLEDGEMENTS

We thank Patrick Sung (UTHSC at San Antonio) for providing yeast RPA and all members of Mazin’s lab for comments and discussion. This work is supported by National Cancer Institute of the National Institutes of Health (NIH) grant numbers CA188347, P30CA056036, and by Drexel-Coulter Program Award (to AM), by National Institute of General Medical Sciences grants R35 GM127085 (to TT), and R01 GM095758 (to FK).

## CONTACT FOR REAGENT AND RESOURCE SHARING

Further information and requests for resources and reagents should be directed to and will be fulfilled by the corresponding author, Dr. Alexander Mazin.

## REFERENCES

Aguilera, A., and Garcia-Muse, T. (2012). R loops: from transcription byproducts to threats to genome stability. Mol. Cell 46, 115–124.

Baranovskiy, A.G., Babayeva, N.D., Suwa, Y., Gu, J., Pavlov, Y.I., and Tahirov, T.H. (2014). Structural basis for inhibition of DNA replication by aphidicolin. Nucleic Acids Res 42, 14013–14021.

Bolger, A.M., Lohse, M., and Usadel, B. (2014). Trimmomatic: a flexible trimmer for Illumina sequence data. Bioinformatics 30, 2114–2120.

Borgstahl, G.E., Brader, K., Mosel, A., Liu, S., Kremmer, E., Goettsch, K.A., Kolar, C., Nasheuer, H.P., and Oakley, G.G. (2014). Interplay of DNA damage and cell cycle signaling at the level of human replication protein A. DNA Repair (Amst) 21, 12–23.

Bugreev, D.V., Hanaoka, F., and Mazin, A.V. (2007). Rad54 dissociates homologous recombination intermediates by branch migration. Nat. Struct. Mol. Biol. 14, 746–753.

Chakraborty, A., Tapryal, N., Venkova, T., Horikoshi, N., Pandita, R.K., Sarker, A.H., Sarkar, P.S., Pandita, T.K., and Hazra, T.K. (2016). Classical non-homologous end-joining pathway utilizes nascent RNA for error-free double-strand break repair of transcribed genes. Nat Commun 7, 13049.

Chedin, F. (2016). Nascent Connections: R-Loops and Chromatin Patterning. Trends Genet 32, 828–838.

Chen, C.C., Lee, J.C., and Chang, M.C. (2012). 4E-BP3 regulates eIF4E-mediated nuclear mRNA export and interacts with replication protein A2. FEBS Lett 586, 2260–2266.

Chen, R., Subramanyam, S., Elcock, A.H., Spies, M., and Wold, M.S. (2016). Dynamic binding of replication protein a is required for DNA repair. Nucleic Acids Res. 44, 5758–5772.

Chen, R., and Wold, M.S. (2014). Replication protein A: single-stranded DNA’s first responder: dynamic DNA-interactions allow replication protein A to direct single-strand DNA intermediates into different pathways for synthesis or repair. BioEssays 36, 1156–1161.

Clewell, D.B., and Helinski, D.R. (1969). Supercoiled circular DNA-protein complex in Escherichia coli: purification and induced conversion to an opern circular DNA form. Proc. Natl. Acad. Sci. USA 62, 1159–1166.

Dobin, A., Davis, C.A., Schlesinger, F., Drenkow, J., Zaleski, C., Jha, S., Batut, P., Chaisson, M., and Gingeras, T.R. (2013). STAR: ultrafast universal RNA-seq aligner. Bioinformatics 29, 15–21.

Fairman, M.P., and Stillman, B. (1988). Cellular factors required for multiple stages of SV40 DNA replication in vitro. EMBO J. 7, 1211–1218.

Gibb, B., Ye, L.F., Gergoudis, S.C., Kwon, Y., Niu, H., Sung, P., and Greene, E.C. (2014). Concentration-dependent exchange of replication protein A on single-stranded DNA revealed by single-molecule imaging. PLoS ONE 9, e87922.

Haring, S.J., Mason, A.C., Binz, S.K., and Wold, M.S. (2008). Cellular functions of human RPA1. Multiple roles of domains in replication, repair, and checkpoints. J. Biol. Chem. 283, 19095–19111.

He, C., Sidoli, S., Warneford-Thomson, R., Tatomer, D.C., Wilusz, J.E., Garcia, B.A., and Bonasio, R. (2016). High-Resolution Mapping of RNA-Binding Regions in the Nuclear Proteome of Embryonic Stem Cells. Mol. Cell 64, 416–430.

Henricksen, L.A., Umbricht, C.B., and Wold, M.S. (1994). Recombinant replication protein A: expression, complex formation, and functional characterization. J. Biol. Chem. 269, 11121–11132.

Itoh, T., and Tomizawa, J. (1980). Formation of an RNA primer for initiation of replication of ColE1 DNA by ribonuclease H. Proc. Natl. Acad. Sci. USA 77, 2450–2454.

Kadyrov, F.A., Genschel, J., Fang, Y., Penland, E., Edelmann, W., and Modrich, P. (2009). A possible mechanism for exonuclease 1-independent eukaryotic mismatch repair. Proc. Natl. Acad. Sci. USA 106, 8495–8500.

Kersey, P.J., Allen, J.E., Armean, I., Boddu, S., Bolt, B.J., Carvalho-Silva, D., Christensen, M., Davis, P., Falin, L.J., Grabmueller, C., et al. (2016). Ensembl Genomes 2016: more genomes, more complexity. Nucleic Acids Res. 44, D574–580.

Keskin, H., Shen, Y., Huang, F., Patel, M., Yang, T., Ashley, K., Mazin, A.V., and Storici, F. (2014). Transcript-RNA-templated DNA recombination and repair. Nature 515, 436–439.

Kim, C., Snyder, R.O., and Wold, M.S. (1992). Binding properties of replication protein A from human and yeast cells. Mol. Cell. Biol. 12, 3050–3059.

Kogoma, T. (1997). Stable DNA replication: interplay between DNA replication, homologous recombination, and transcription. Microbiol. Mol. Biol. Rev. 61, 212–238.

Laemmli, U.K. (1970). Cleavage of structural proteins during the assembly of the head of bacteriophage T4. Nature 227, 680–685.

Lang, K.S., Hall, A.N., Merrikh, C.N., Ragheb, M., Tabakh, H., Pollock, A.J., Woodward, J.J., Dreifus, J.E., and Merrikh, H. (2017). Replication-Transcription Conflicts Generate R-Loops that Orchestrate Bacterial Stress Survival and Pathogenesis. Cell 170, 787–799 e718.

Lee, S., Kopp, F., Chang, T.C., Sataluri, A., Chen, B., Sivakumar, S., Yu, H., Xie, Y., and Mendell, J.T. (2016). Noncoding RNA NORAD Regulates Genomic Stability by Sequestering PUMILIO Proteins. Cell 164, 69–80.

Li, H., Handsaker, B., Wysoker, A., Fennell, T., Ruan, J., Homer, N., Marth, G., Abecasis, G., Durbin, R., and Genome Project Data Processing, S. (2009). The Sequence Alignment/Map format and SAMtools. Bioinformatics 25, 2078–2079.

Marnef, A., Cohen, S., and Legube, G. (2017). Transcription-Coupled DNA Double-Strand Break Repair: Active Genes Need Special Care. J. Mol. Biol. 429, 1277–1288.

Mass, G., Nethanel, T., and Kaufmann, G. (1998). The middle subunit of replication protein A contacts growing RNA-DNA primers in replicating simian virus 40 chromosomes. Mol. Cell. Biol. 18, 6399–6407.

Masutani, C., Kusumoto, R., Iwai, S., and Hanaoka, F. (2000). Mechanisms of accurate translesion synthesis by human DNA polymerase eta. EMBO J. 19, 3100–3109.

Mazina, O.M., Keskin, H., Hanamshet, K., Storici, F., and Mazin, A.V. (2017). Rad52 Inverse Strand Exchange Drives RNA-Templated DNA Double-Strand Break Repair. Mol. Cell 67, 19–29 e13.

McIlwraith, M.J., Vaisman, A., Liu, Y., Fanning, E., Woodgate, R., and West, S.C. (2005). Human DNA polymerase eta promotes DNA synthesis from strand invasion intermediates of homologous recombination. Mol. Cell 20, 783–792.

Mitchell, S.F., Jain, S., She, M., and Parker, R. (2013). Global analysis of yeast mRNPs. Nat. Struct. Mol. Biol. 20, 127–133.

Mo, J., Liu, L., Leon, A., Mazloum, N., and Lee, M.Y. (2000). Evidence that DNA polymerase delta isolated by immunoaffinity chromatography exhibits high-molecular weight characteristics and is associated with the KIAA0039 protein and RPA. Biochemistry 39, 7245–7254.

Nguyen, B., Sokoloski, J., Galletto, R., Elson, E.L., Wold, M.S., and Lohman, T.M. (2014). Diffusion of human replication protein A along single-stranded DNA. J. Mol. Biol. 426, 3246–3261.

Nguyen, H.D., Yadav, T., Giri, S., Saez, B., Graubert, T.A., and Zou, L. (2017). Functions of Replication Protein A as a Sensor of R Loops and a Regulator of RNaseH1. Mol. Cell 65, 832–847 e834.

Plosky, B.S., and Woodgate, R. (2004). Switching from high-fidelity replicases to low-fidelity lesion-bypass polymerases. Curr Opin Genet Dev 14, 113–119.

Pokhrel, N., Caldwell, C.C., Corless, E.I., Tillison, E.A., Tibbs, J., Jocic, N., Tabei, S.M.A., Wold, M.S., Spies, M., and Antony, E. (2019). Dynamics and selective remodeling of the DNA-binding domains of RPA. Nat. Struct. Mol. Biol. 26, 129–136.

Rodriges Blanko, E., Kadyrova, L.Y., and Kadyrov, F.A. (2016). DNA Mismatch Repair Interacts with CAF-1- and ASF1A-H3-H4-dependent Histone (H3-H4)2 Tetramer Deposition. J. Biol. Chem. 291, 9203–9217.

Rossi, M.J., Mazina, O.M., Bugreev, D.V., and Mazin, A.V. (2010). Analyzing the branch migration activities of eukaryotic proteins. Methods 51, 336–346.

Santos-Pereira, J.M., and Aguilera, A. (2015). R loops: new modulators of genome dynamics and function. Nat Rev Genet 16, 583–597.

Sanz, L.A., Hartono, S.R., Lim, Y.W., Steyaert, S., Rajpurkar, A., Ginno, P.A., Xu, X., and Chedin, F. (2016). Prevalent, Dynamic, and Conserved R-Loop Structures Associate with Specific Epigenomic Signatures in Mammals. Mol. Cell 63, 167–178.

Sigurdsson, S., Trujillo, K., Song, B., Stratton, S., and Sung, P. (2001). Basis for avid homologous DNA strand exchange by human Rad51 and RPA. J. Biol. Chem. 276, 8798–8806.

Singleton, M.R., Wentzell, L.M., Liu, Y., West, S.C., and Wigley, D.B. (2002). Structure of the single-strand annealing domain of human RAD52 protein. Proc. Natl. Acad. Sci. USA 99, 13492–13497.

Song, B., and Sung, P. (2000). Functional interactions among yeast Rad51 recombinase, Rad52 mediator, and replication protein A in DNA strand exchange. J. Biol. Chem. 275, 15895–15904.

Stith, C.M., Sterling, J., Resnick, M.A., Gordenin, D.A., and Burgers, P.M. (2008). Flexibility of eukaryotic Okazaki fragment maturation through regulated strand displacement synthesis. J. Biol. Chem. 283, 34129–34140.

Stork, C.T., Bocek, M., Crossley, M.P., Sollier, J., Sanz, L.A., Chedin, F., Swigut, T., and Cimprich, K.A. (2016). Co-transcriptional R-loops are the main cause of estrogen-induced DNA damage. Elife 5.

Stuckey, R., Garcia-Rodriguez, N., Aguilera, A., and Wellinger, R.E. (2015). Role for RNA:DNA hybrids in origin-independent replication priming in a eukaryotic system. Proc. Natl. Acad. Sci. USA 112, 5779–5784.

Trapnell, C., Roberts, A., Goff, L., Pertea, G., Kim, D., Kelley, D.R., Pimentel, H., Salzberg, S.L., Rinn, J.L., and Pachter, L. (2012). Differential gene and transcript expression analysis of RNA-seq experiments with TopHat and Cufflinks. Nat Protoc 7, 562–578.

Tresini, M., Warmerdam, D.O., Kolovos, P., Snijder, L., Vrouwe, M.G., Demmers, J.A., van, I.W.F., Grosveld, F.G., Medema, R.H., Hoeijmakers, J.H., et al. (2015). The core spliceosome as target and effector of non-canonical ATM signalling. Nature 523, 53–58.

Wei, L., Levine, A.S., and Lan, L. (2016). Transcription-coupled homologous recombination after oxidative damage. DNA Repair (Amst) 44, 76–80.

Wei, L., Nakajima, S., Bohm, S., Bernstein, K.A., Shen, Z., Tsang, M., Levine, A.S., and Lan, L. (2015). DNA damage during the G0/G1 phase triggers RNA-templated, Cockayne syndrome B-dependent homologous recombination. Proc. Natl. Acad. Sci. USA 112, E3495–3504.

Wobbe, C.R., Weissbach, L., Borowiec, J.A., Dean, F.B., Murakami, Y., Bullock, P., and Hurwitz, J. (1987). Replication of simian virus 40 origin-containing DNA in vitro with purified proteins. Proc. Natl. Acad. Sci. USA 84, 1834–1838.

Wold, M.S. (1997). Replication protein A: a heterotrimeric, single-stranded DNA-binding protein required for eukaryotic DNA metabolism. Ann. Rev. Biochem. 66, 61–92.

Wold, M.S., and Kelly, T. (1988). Purification and characterization of replication protein A, a cellular protein required for in vitro replication of simian virus 40 DNA. Proc. Natl. Acad. Sci. USA 85, 2523–2527.

Yasuhara, T., Kato, R., Hagiwara, Y., Shiotani, B., Yamauchi, M., Nakada, S., Shibata, A., and Miyagawa, K. (2018). Human Rad52 Promotes XPG-Mediated R-loop Processing to Initiate Transcription-Associated Homologous Recombination Repair. Cell 175, 558–570 e511.

Yuzhakov, A., Kelman, Z., Hurwitz, J., and O’Donnell, M. (1999). Multiple competition reactions for RPA order the assembly of the DNA polymerase delta holoenzyme. EMBO J. 18, 6189–6199.

Zahurancik, W.J., Klein, S.J., and Suo, Z. (2013). Kinetic mechanism of DNA polymerization catalyzed by human DNA polymerase epsilon. Biochemistry 52, 7041–7049.

Zaitsev, E.N., and Kowalczykowski, S.C. (2000). A novel pairing process promoted by Escherichia coli RecA protein: inverse DNA and RNA strand exchange. Genes Dev. 14, 740–749.

Zou, Y., Liu, Y., Wu, X., and Shell, S.M. (2006). Functions of human replication protein A (RPA): from DNA replication to DNA damage and stress responses. J Cell Physiol 208, 267–273.

