## Supplementary figures S1-S11, Supplementary Table S1-S2. for "Replication protein A binds RNA and promotes R-loop formation"

**CONTENTS:**

1. Supplementary Figures 1-11
2. Supplementary Tables 1-2

Fig. S1

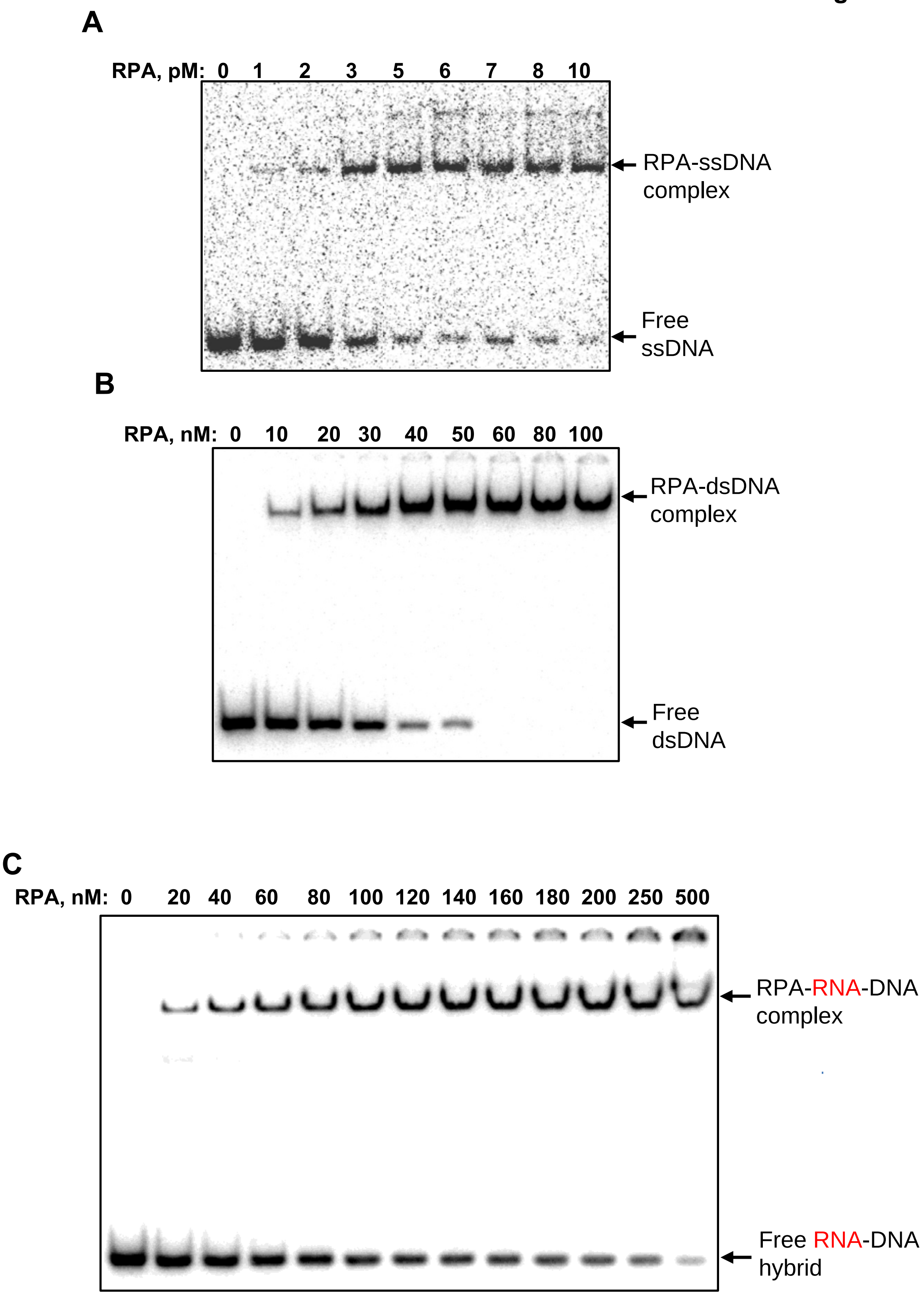
A
RPA, pM: 0 1 2 3 5 6 7 8 10
RPA-ssDNA complex
Free ssDNA
B
RPA, nM: 0 10 20 30 40 50 60 80 100
RPA-dsDNA complex
Free dsDNA
C
RPA, nM: 0 20 40 60 80 100 120 140 160 180 200 250 500
RPA-RNA-DNA complex
Free RNA-DNA hybrid

**Figure S1. Related to Figure 1. Binding affinity of human RPA for ssDNA, dsDNA and RNA-DNA hybrid.** Analysis of human RPA binding to: **A,** $^{32}$P-labeled ssDNA (48-nt, no.211, 0.5 pM), **B,** to $^{32}$P-labeled dsDNA (48-bp no.211/no.212, 3 nM) and **C,** to $^{32}$P-labeled RNA-DNA hybrid (48-bp no.501/no.212, 3 nM) using EMSA in a 6% polyacrylamide gel. The RPA concentrations are indicated at the top of the gels.

**Fig. S2**

**A**

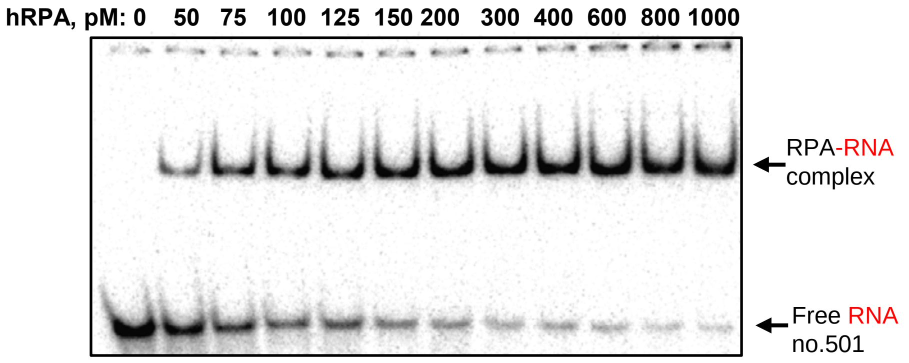
hRPA, pM: 0 50 75 100 125 150 200 300 400 600 800 1000
RPA-RNA complex
Free RNA no.501

**B**

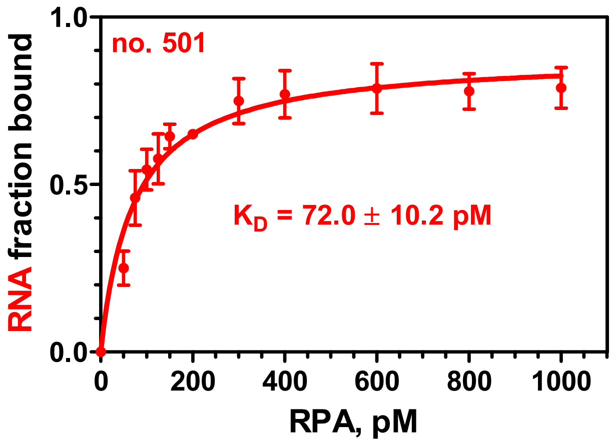
no. 501
K_D = 72.0 ± 10.2 pM
RNA fraction bound
RPA, pM
1.0
0.5
0.0
0
200
400
600
800
1000

**Figure S2. Related to Figure 1. RPA binding to RNA in the presence of 100 mM NaCl. A,** Analysis of RPA binding to a 48-mer RNA (no. 501; 5 pM) using EMSA in a 6% polyacrylamide gel. **B,** Data from (A) plotted as a graph. The $K_D$ values were obtained by fitting the data to one site binding hyperbola in GraphPad Prism 5.0. The error bars indicate standard error of the mean (S.E.).

Fig. S3

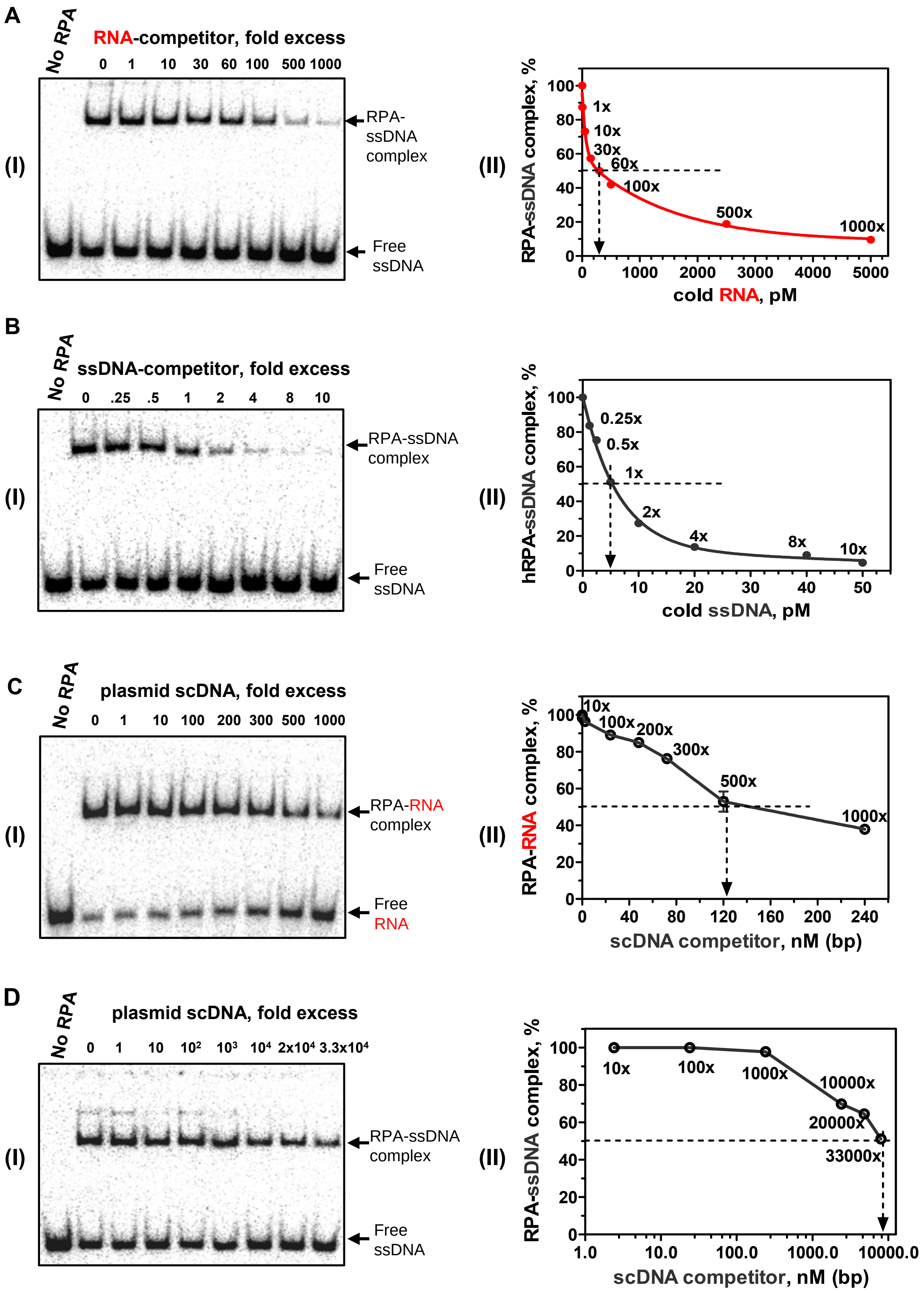
A
No RPA
RNA-competitor, fold excess
0 1 10 30 60 100 500 1000
RPA-ssDNA complex
Free ssDNA
(I)
(II)
RPA-ssDNA complex, %
100 80 60 40 20 0
1x
10x
30x
60x
100x
500x
1000x
0 1000 2000 3000 4000 5000
cold RNA, pM
B
No RPA
ssDNA-competitor, fold excess
0 .25 .5 1 2 4 8 10
RPA-ssDNA complex
Free ssDNA
(I)
(II)
hRPA-ssDNA complex, %
100 80 60 40 20 0
0.25x
0.5x
1x
2x
4x
8x
10x
0 10 20 30 40 50
cold ssDNA, pM
C
No RPA
plasmid scDNA, fold excess
0 1 10 100 200 300 500 1000
RPA-RNA complex
Free RNA
(I)
(II)
RPA-RNA complex, %
100 80 60 40 20 0
10x
100x
200x
300x
500x
1000x
0 40 80 120 160 200 240
scDNA competitor, nM (bp)
D
No RPA
plasmid scDNA, fold excess
0 1 10 10² 10³ 10⁴ 2x10⁴ 3.3x10⁴
RPA-ssDNA complex
Free ssDNA
(I)
(II)
RPA-ssDNA complex, %
100 80 60 40 20 0
10x
100x
1000x
10000x
20000x
33000x
1.0 10.0 100.0 1000.0 10000.0
scDNA competitor, nM (bp)

**Figure S3. Related to Figure 1. Analysis of RPA binding to DNA and RNA using RNA- and DNA-competitors.** Analysis of human RPA (20 pM) binding to $^{32}P$-labeled ssDNA (no.211, 5 pM) in the presence of increasing amounts of non-labeled identical RNA (no. 501; 0, 5, 50, 150, 300, 500, 2500, and 5000 pM) (**A,** panel **I**); or non-labeled identical ssDNA (no. 211, 0, 1.25, 2.5, 5, 10, 20, 40, and 50 pM) (**B,** panel **I**) using EMSA in a 6% polyacrylamide gel. The molar excesses of RNA or cold ssDNA over $^{32}P$-labeled ssDNA are shown above the gels. (**A-B**, panels **II**) graphical representation of data from (**A-B,** panels **I**), respectively. RPA binding to ssDNA (5 pM) was reduced by 50% (indicated by the dashed lines) in the presence of 300 pM (60 x) of RNA or 5 pM (1 x) of non-labeled ssDNA. (**C**, panel **I**) RPA (125 pM) binding to $^{32}P$-labeled RNA (no.501, 5 pM or 0.24 nM, nt) in the presence of increasing amounts of non-homologous supercoiled (sc) pHSG299 plasmid dsDNA (0, 0.24, 2.4, 24, 48, 72, 120, and 240 nM, bp), or (**D,** panel **I**) RPA (30 pM) binding to $^{32}P$-labeled ssDNA (no.211, 5 pM or 0.24 nM, nt) in the presence of pHSG299 plasmid scDNA (0, 0.24, 2.4, 24, 240, 2400, 4800, and 8000 nM, bp) using EMSA in a 6% polyacrylamide gel. Molar excesses of dsDNA (in bp) over RNA or ssDNA (in nt) are shown above the gels. (**C-D**, panels **II**) Graphical representation of the data from of (**C-D**, panels **I**), respectively. RPA binding to RNA (0.24 nM, nt) or ssDNA (0.24 nM, nt) was reduced by 50% (indicated by the dashed lines) in the presence of 120 nM (bp) (500 x) or 8000 nM (bp) (33,000 x) of supercoiled plasmid dsDNA. The error bars indicate S.E.

Fig. S4

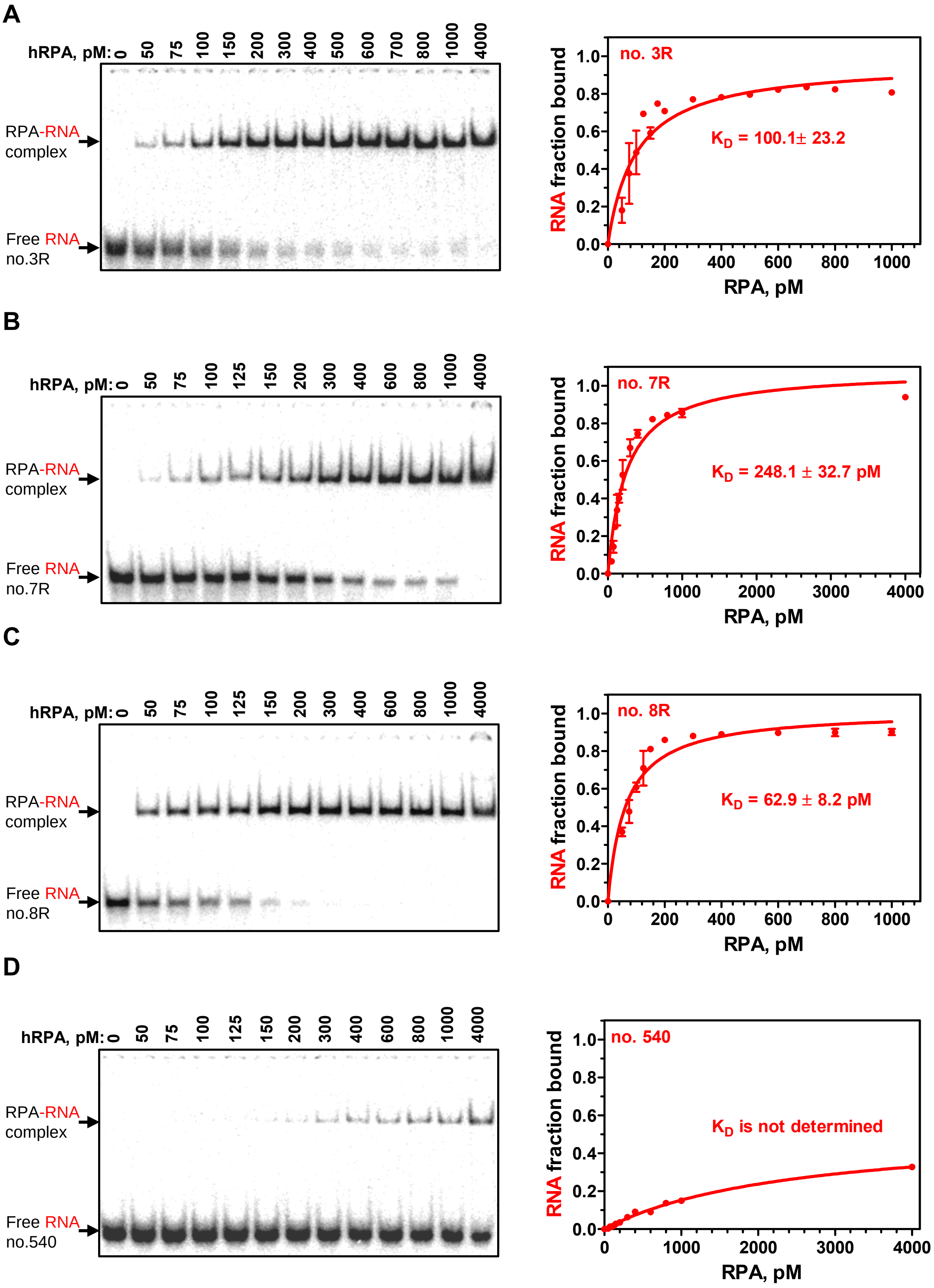

A
hRPA, pM: 0 50 75 100 150 200 300 400 500 600 700 800 1000 4000
RPA-RNA complex
Free RNA no.3R
no. 3R
$K_D$ = 100.1± 23.2
RNA fraction bound
RPA, pM
B
hRPA, pM: 0 50 75 100 125 150 200 300 400 600 800 1000 4000
RPA-RNA complex
Free RNA no.7R
no. 7R
$K_D$ = 248.1 ± 32.7 pM
RNA fraction bound
RPA, pM
C
hRPA, pM: 0 50 75 100 125 150 200 300 400 600 800 1000 4000
RPA-RNA complex
Free RNA no.8R
no. 8R
$K_D$ = 62.9 ± 8.2 pM
RNA fraction bound
RPA, pM
D
hRPA, pM: 0 50 75 100 125 150 200 300 400 600 800 1000 4000
RPA-RNA complex
Free RNA no.540
no. 540
$K_D$ is not determined
RNA fraction bound
RPA, pM

**Figure S4. Related to Figure 1. RPA binding to RNAs of different sequences.** Analysis of RPA binding to a 48-mer RNAs (5 pM) (no. 3R) (**A**), (no.7R) (**B**), (no.8R) (**C**) and (no. 540) (**D**) using EMSA in a 6% polyacrylamide gel. On right panels, the data from (A-D) are plotted as graphs. The $K_D$ values were obtained by fitting the data to one site binding hyperbola in GraphPad Prism 5.0. The error bars indicate standard error of the mean (S.E.).

**Figure S5**

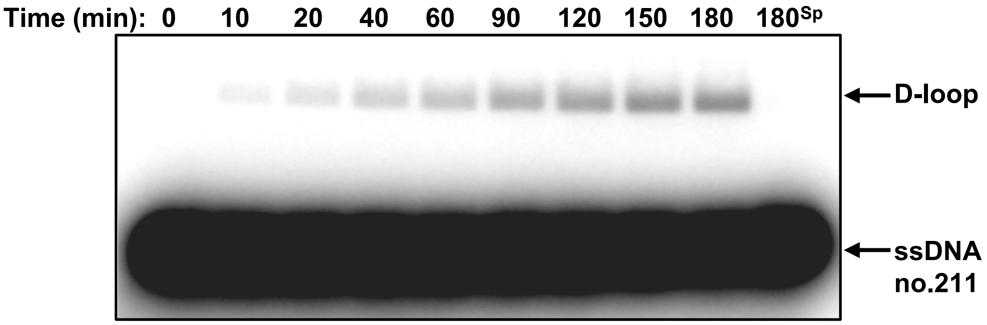

**Figure S5. Related to Figure 2. Human RPA promotes D-loop formation with low efficiency.** RPA (200 nM) was preincubated with a 48-mer ssDNA (no.211; 3 μM, nt) for 15 min. D-loop formation was initiated by addition of supercoiled pUC19 dsDNA (67.2 μM, nt) and carried out for the indicated periods of time. The reaction products were analyzed by electrophoresis in a 1% agarose gel. “$180^{sp}$” denotes the RPA-independent (spontaneous) D-loop formation after 180 min of reaction. The quantification of the gel is shown in Figure 2C.

**Fig. S6**

**A**

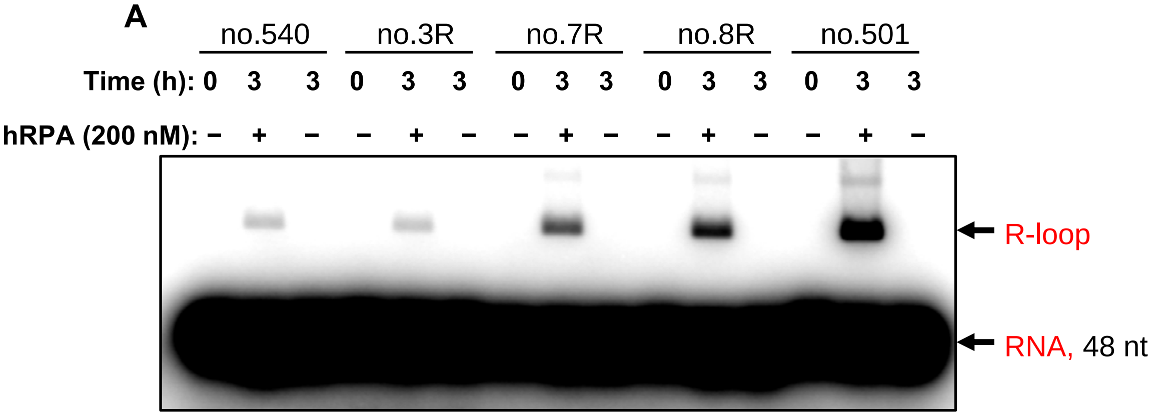

no.540
no.3R
no.7R
no.8R
no.501
Time (h): 0 3 3 0 3 3 0 3 3 0 3 3 0 3 3
hRPA (200 nM): – + – – + – – + – – + – – + –
R-loop
RNA, 48 nt


**B**

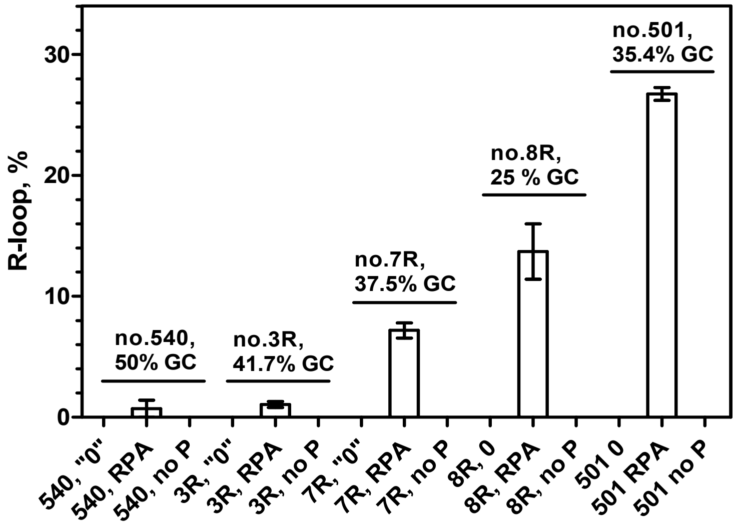

R-loop, %
0
10
20
30
no.540, 50% GC
no.3R, 41.7% GC
no.7R, 37.5% GC
no.8R, 25 % GC
no.501, 35.4% GC
540, "0"
540, RPA
540, no P
3R, "0"
3R, RPA
3R, no P
7R, "0"
7R, RPA
7R, no P
8R, 0
8R, RPA
8R, no P
501 0
501 RPA
501 no P

**Figure S6. Related to Figure 2. RNA-sequence dependence of R-loop formation by RPA. A**, Five 48-mer RNAs of different sequences (each 3 μM, nt) were tested in R-loop formation reaction promoted by RPA. The experiments were performed as in Figure 2. The R-loops were analyzed by electrophoresis in a 1% agarose gel. **B,** The data from panel A presented as a graph. Percentage of GC pairs (%GC) for each RNA substrate is shown above the graph. The error bars indicate S.E.

Fig. S7

A

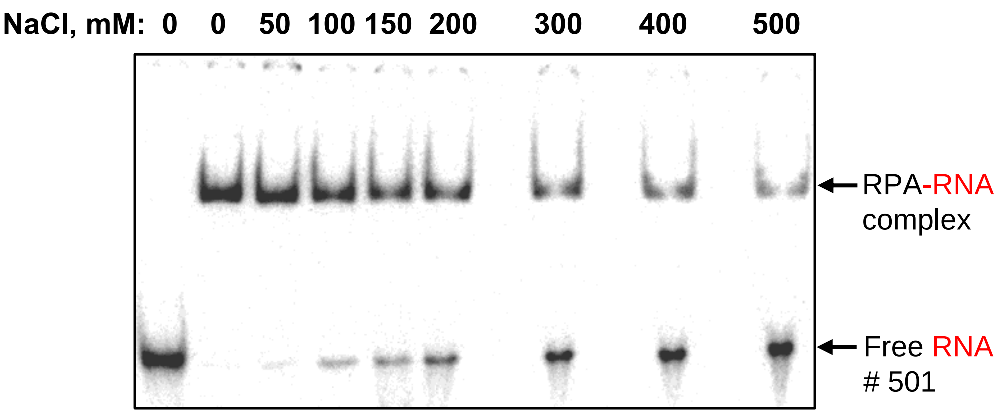
NaCl, mM: 0 0 50 100 150 200 300 400 500
RPA-RNA complex
Free RNA # 501

B

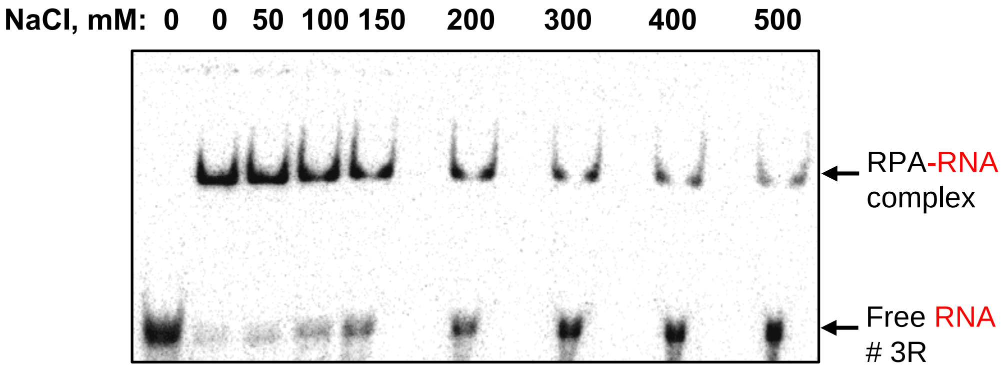
NaCl, mM: 0 0 50 100 150 200 300 400 500
RPA-RNA complex
Free RNA # 3R

C

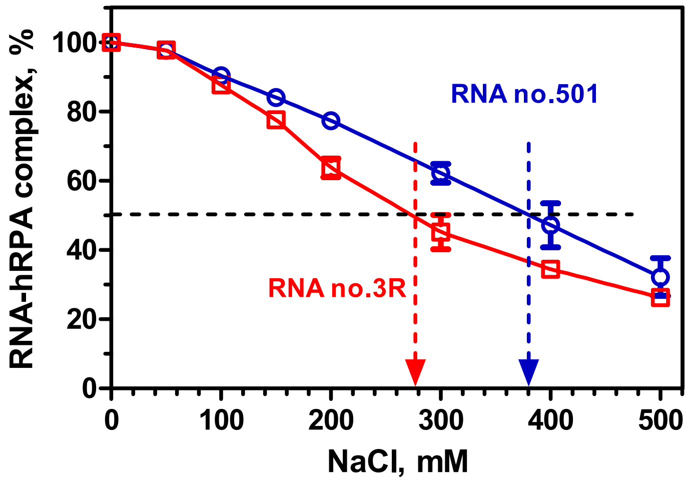
RNA-hRPA complex, %
100
80
60
40
20
0
0
100
200
300
400
500
NaCl, mM
RNA no.501
RNA no.3R

**Figure S7. Related to Figure 2. The effect of NaCl concentration on the stability of the RPA-RNA complexes.**

**A, B,** RPA (200 pM) was incubated with RNA molecules (no.501; 5 pM) or (no.3R; 5 pM) for 15 min at 37 °C in binding buffer supplemented with NaCl in the indicated concentrations. The samples were analyzed using EMSA in 6% polyacrylamide gels. **C,** The data from A and B presented as a graph. The vertical arrows indicate the salt titration midpoint of the RPA-RNA complexes, which were 380 mM and 280 mM for RPA-RNA complexes formed with RNA no.501 and no.3R, respectively.

Fig. S8

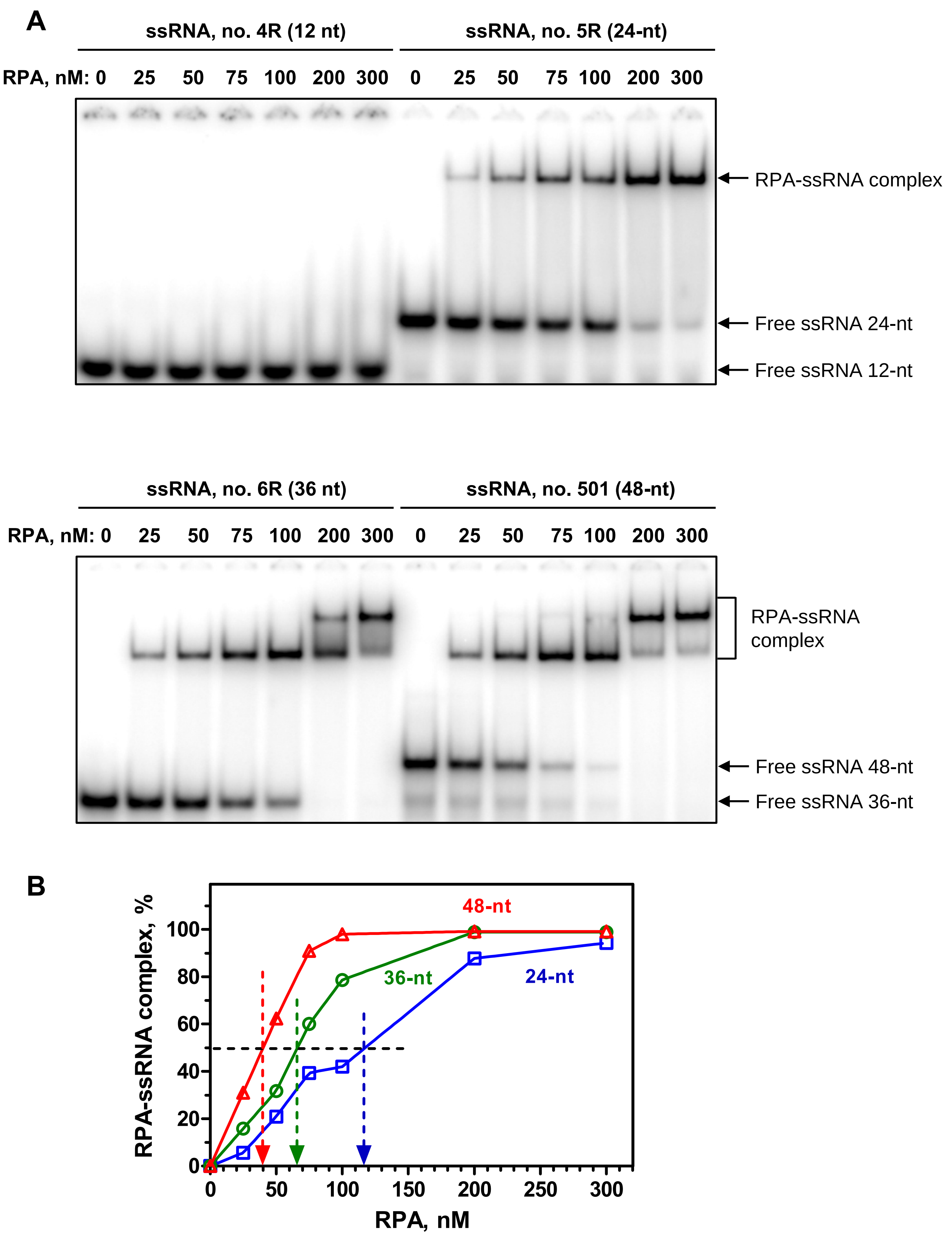

A
ssRNA, no. 4R (12 nt)
ssRNA, no. 5R (24-nt)
RPA, nM: 0 25 50 75 100 200 300 0 25 50 75 100 200 300
RPA-ssRNA complex
Free ssRNA 24-nt
Free ssRNA 12-nt
ssRNA, no. 6R (36 nt)
ssRNA, no. 501 (48-nt)
RPA, nM: 0 25 50 75 100 200 300 0 25 50 75 100 200 300
RPA-ssRNA complex
Free ssRNA 48-nt
Free ssRNA 36-nt
B
RPA-ssRNA complex, %
RPA, nM
48-nt
36-nt
24-nt
0
20
40
60
80
100
0
50
100
150
200
250
300

**Figure S8. Related to Figure 2.   Related to Figure 2. RPA binds RNA in a length dependent manner. A,** Binding of RPA (at indicated concentrations) to a 12-, 24-, 36-, and 48-nt long RNA (each 3 μM, nt) under R-loop formation conditions for 15 min at 37 °C was examined using EMSA. **B,** The data from panel A presented as a graph. The protein concentrations that correspond to 50% of RNA binding is indicated by the dashed lines, and were 40, 65, and 120 nM for 48-nt, 36-nt, and 24-nt long RNA, respectively.

Fig. S9

A

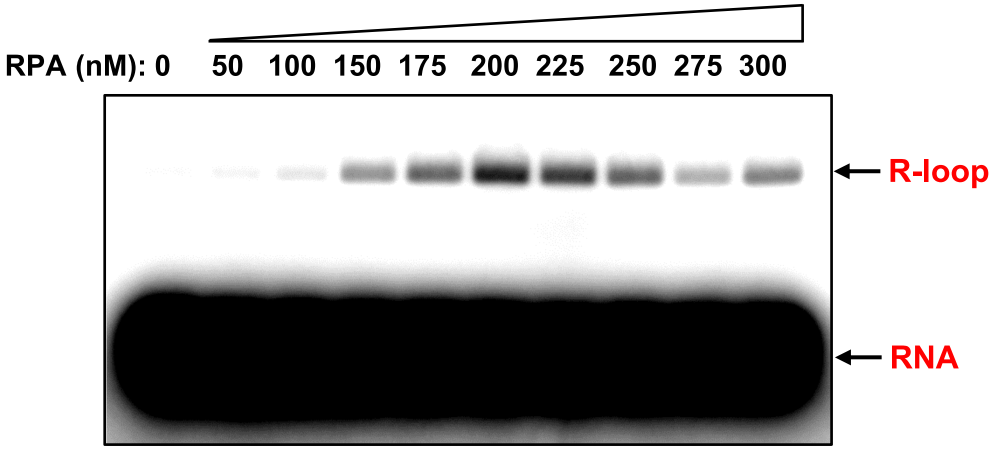
RPA (nM): 0
50
100
150
175
200
225
250
275
300
R-loop
RNA

B

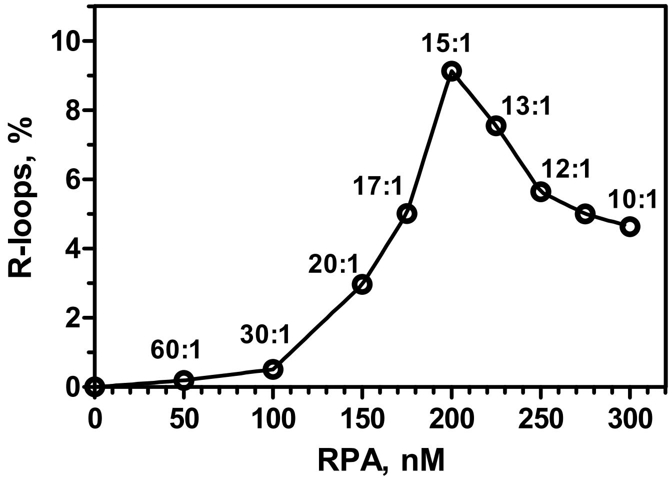
R-loops, %
RPA, nM
0
2
4
6
8
10
50
100
150
200
250
300
60:1
30:1
20:1
17:1
15:1
13:1
12:1
10:1

**Figure S9. Related to Figure 2. Effect of RPA concentration on the R-loop formation efficiency. A,** RPA in indicated concentrations was incubated with RNA (no.501; 3 μM, nt), and R-loop formation was initiated by addition of supercoiled pUC19 dsDNA. The reactions were carried out for 30 min at 37 °C. **B,** The data from A shown as a graph. The ratios of RNA nucleotides per one RPA trimer are shown above the curve.

Fig S10

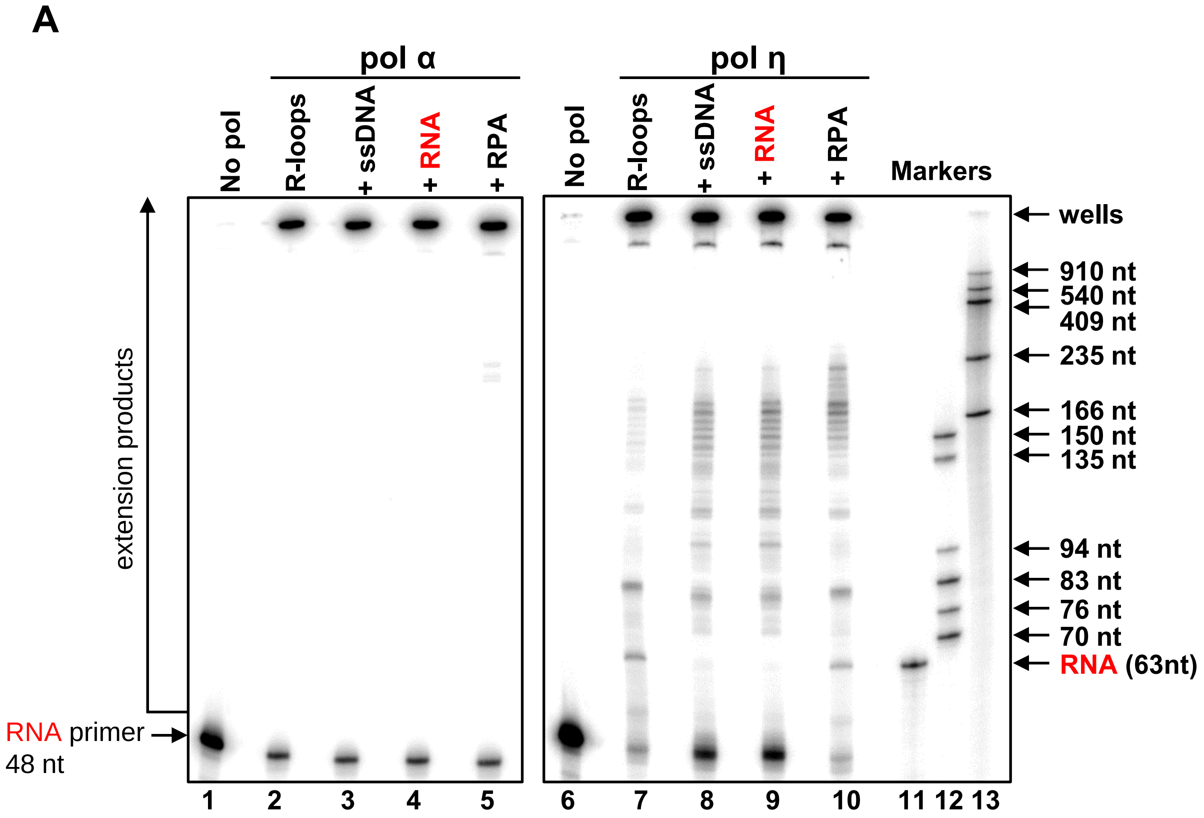

A
pol α
pol η
No pol
R-loops
+ ssDNA
+ RNA
+ RPA
Markers
wells
910 nt
540 nt
409 nt
235 nt
166 nt
150 nt
135 nt
94 nt
83 nt
76 nt
70 nt
RNA (63nt)
extension products
RNA primer
48 nt
1
2
3
4
5
6
7
8
9
10
11 12 13


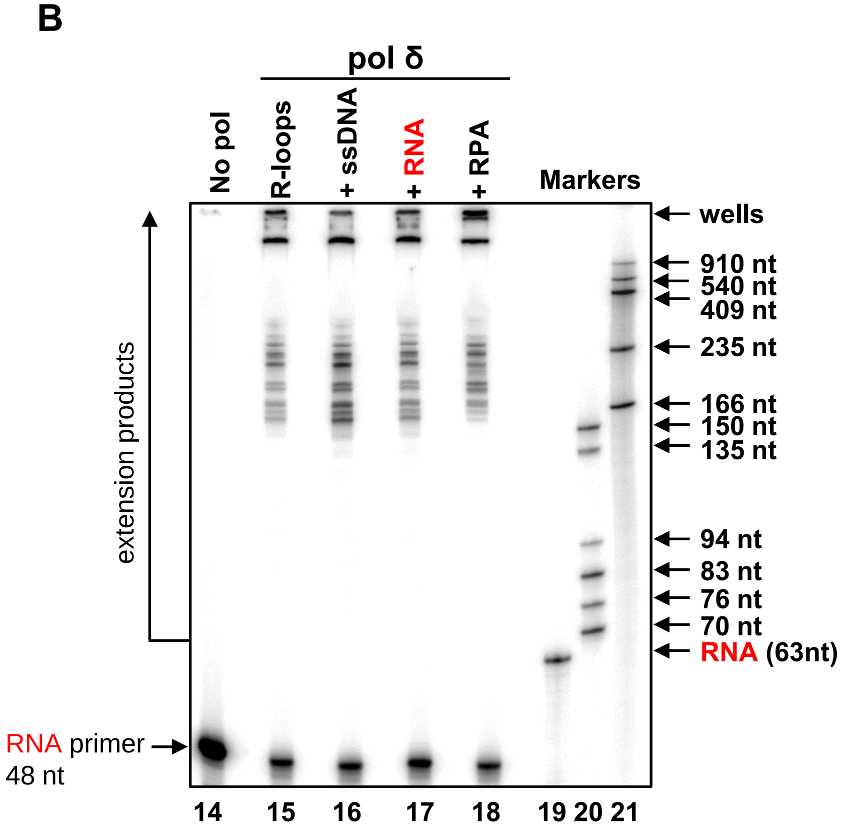

B
pol δ
No pol
R-loops
+ ssDNA
+ RNA
+ RPA
Markers
wells
910 nt
540 nt
409 nt
235 nt
166 nt
150 nt
135 nt
94 nt
83 nt
76 nt
70 nt
RNA (63nt)
extension products
RNA primer
48 nt
14
15
16
17
18
19 20 21

**Figure S10. Related to Figure 3. Effect of RNA, ssDNA and RPA on R-loop mediated DNA synthesis. A, B,** Using deproteinized and purified R-loops (1 nM) RNA extension reactions were carried out using pol α (50 nM), pol η (4 nM), or pol δ (0.5 nM). The reactions that were supplemented with ssDNA (no.2; 3 μM, nt), RNA (no.517; 3 μM, nt), or RPA (5 nM) are indicated above the gels. In controls (lanes 1, 6 and 14) DNA polymerases were substituted by their storage buffers. $^{32}$P-labeled RNA/DNA markers are showed in lanes 11-13 and 19-21.

Fig. S11

A

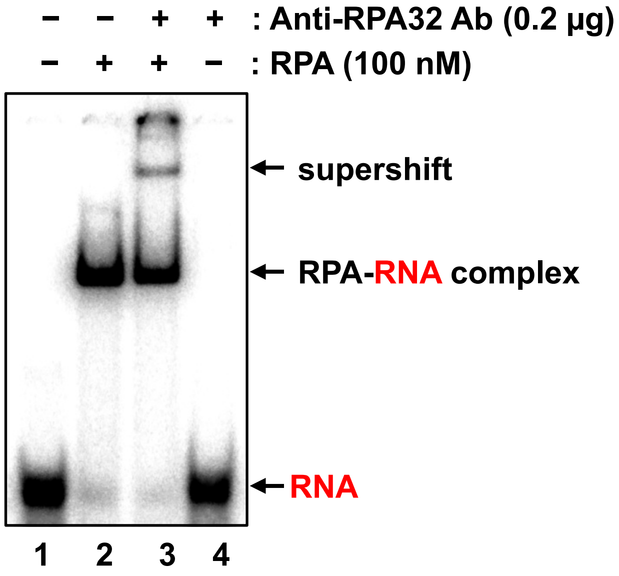

– – + + : Anti-RPA32 Ab (0.2 μg)
– + + – : RPA (100 nM)
supershift
RPA-RNA complex
RNA
1 2 3 4


B

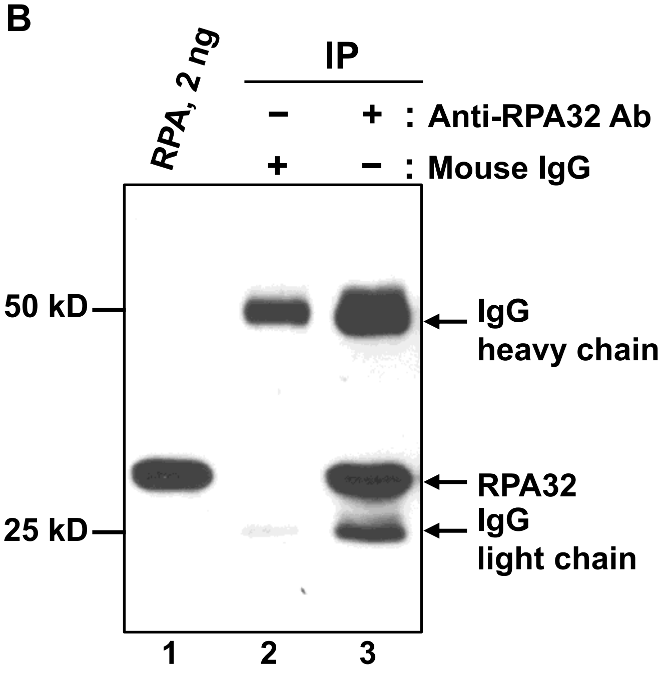

RPA, 2 ng
IP
– + : Anti-RPA32 Ab
+ – : Mouse IgG
50 kD
25 kD
IgG
heavy chain
RPA32
IgG
light chain
1 2 3

**Figure S11. Related to Figure 6. Characterization of the RPA-RNA complexes using primary antibody against human RPA32.** (**A**) Primary anti-RPA32 antibodies can efficiently interact with RPA-RNA complex. $^{32}$P-labeled RNA (3 μM, nt) was incubated in the absence (lane 1) or presence RPA (100 nM) (lane 2) for 15 min at 37 °C. Primary mouse polyclonal anti-RPA32 antibodies (0.2 μg, ab88675) were added to the pre-formed RPA-RNA complex (lane 3) or to $^{32}$P- labeled RNA (lane 4) followed by an 1-h incubation on ice. The samples were analyzed by a 6% native polyacrylamide (29:1) gel. (**B**) Detection of RPA in the immunoprecipitants from HEK 293T cells. Whole cell extracts (WCE) (750 μg) from HEK 293T cells were pre-cleared with 3 μg of normal mouse immunoglobulin G (IgG) (lane 2) and then RPA was immunoprecipitated (IP) using 3 μg of anti-RPA32 specific antibody (lane3). The immunoprecipitated protein samples (2.5% of total) were resolved by a 10% SDS-PAGE, transferred to PDVF membranes, and probed with anti-RPA32 antibodies (ab88675). Purified RPA (2 ng) was used as a marker (lane 1).

**Table S1. Sequences of the oligonucleotides used in this study**

| Number | Length, nt | DNA/ RNA | Sequence (5'→3') |
|---|---|---|---|
| 2 | 63 | DNA | TCCTTTTGATAAGAGGTCATTTTTGCGGATGGCTTAGAGC TTAATTGCTGAATCTGGTGCTGT |
| 211 | 48 | DNA | GAAGCATTTATCAGGGTTATTGTCTCATGAGCGGATACAT ATTTGAAT |
| 212 | 48 | DNA | ATTCAAATATGTATCCGCTCATGAGACAATAACCCTGATAA ATGCTTC |
| 501 | 48 | RNA | GAAGCAUUUAUCAGGGUUAUUGUCUCAUGAGCGGAUAC AUAUUUGAAU |
| 517 | 63 | RNA | UCCUUUUGAUAAGAGGUCAUUUUUGCGGAUGGCUUAGA GCUUAAUUGCUGAAUCUGGUGCUGU |
| 540 | 48 | RNA | UGUACUGAGAGUGCACCAUAUGCGGUGUGAAAUACCGC ACAGAUGCGU |
| 3R | 48 | RNA | AAUGCUUAAUCAGUGAGGCACCUAUCUCAGCGAUCUGU CUAUUUCGUU |
| 4R | 12 | RNA | GAUACAUAUUUG |
| 5R | 24 | RNA | GAAGCAUUUAUCAGGGUUAUUGUC |
| 6R | 36 | RNA | GAAGCAUUUAUCAGGGUUAUUGUCUCAUGAGCGGAU |
| 7R | 48 | RNA | CGUUAAGGGAUUUUGGUCAUGAGAUUAUCAAAAAGGAU CUUCACCUAG |
| 8R | 48 | RNA | UUUUAAAUCAAUCUAAAGUAUAUAUGAGUAAACUUGGUC UGACAGUUA |

**Table S2. The minimum free energy of RNA molecules used in this study.**

| RNA no. | Minimum Free Energy, kcal/mol | The R-loop yield, % | $K_D$ of RPA binding to RNA, pM |
|---|---|---|---|
| 501 | -3.5 | 26.7 ± 0.5 | 101.4 ± 17 |
| 8R | -3.8 | 13.7 ± 2.3 | 62.9 ± 8.2 |
| 7R | -5.4 | 7.2 ± 0.6 | 248.1 ± 32.7 |
| 3R | -5.9 | 1.1 ± 0.3 | 100 ± 23.2 |
| 540 | -14.3 | 0.7 ± 0.7 | > 4000 |

The RNA minimum free energy was calculated using ViennaRNA Package 2.0 (*1*).

The data on the R-loop yield are from Figure S6.

### REFERENCE


1. R. Lorenz *et al.*, ViennaRNA Package 2.0. *Algorithms Mol Biol* **6**, 26 (2011).
